# Capsular Polysaccharide Restrains Type VI Secretion in *Acinetobacter baumannii*

**DOI:** 10.1101/2024.04.23.590769

**Authors:** Nicolas Flaugnatti, Loriane Bader, Mary Croisier-Coeytaux, Melanie Blokesch

## Abstract

The type VI secretion system (T6SS) is a sophisticated, contact-dependent nanomachine involved in interbacterial competition. To function effectively, the T6SS must penetrate the membranes of both attacker and target bacteria. Structures associated with the cell envelope, like polysaccharides chains, can therefore introduce spatial separation and steric hindrance, potentially affecting the efficacy of the T6SS. In this study, we examined how the capsular polysaccharide (CPS) of *Acinetobacter baumannii* affects T6SS’s antibacterial function. Our findings show that the CPS confers resistance against T6SS-mediated assaults from rival bacteria. Notably, under typical growth conditions, the presence of the surface-bound capsule also reduces the efficacy of the bacterium’s own T6SS. This T6SS impairment is further enhanced when CPS is overproduced due to genetic modifications or antibiotic treatment. Furthermore, we demonstrate that the bacterium adjusts the level of the T6SS inner tube protein Hcp according to its secretion capacity, by initiating a degradation process involving the ClpXP protease. Collectively, our findings contribute to a better understanding of the dynamic relationship between T6SS and CPS and how they respond swiftly to environmental challenges.

## Introduction

*Acinetobacter baumannii* is an opportunistic pathogen known for causing hospital-acquired infections. The World Health Organization (WHO) has identified it as a critically high-priority pathogen in dire need of new therapeutic strategies (Tacconelli *et al*, 2018). Consistent with its classification, *A. baumannii* is a member of the “ESKAPE bugs”, a term referring to six pathogenic species (***E****nterococcus faecium*, ***S****taphylococcus aureus*, ***K****lebsiella pneumoniae*, ***A****cinetobacter baumannii*, ***P****seudomonas aeruginosa*, ***E****nterobacter spp*.) notorious for causing hospital-acquired infections and their ability to escape antibiotic treatments (Rice, 2008).

*A. baumannii* can gain new functions, including antibiotic resistance, through horizontal gene transfer, notably via plasmid conjugation (Di Venanzio *et al*, 2019; Hamidian *et al*, 2014) and natural competence for transformation (Godeux *et al*, 2018; Godeux *et al*, 2022; Harding *et al*, 2013; Ramirez *et al*, 2010; Vesel & Blokesch, 2021; Wilharm *et al*, 2013). Beyond its resistance to antibiotics, the bacterium can withstand desiccation, disinfectants, and survive on surfaces for extended periods, posing a significant challenge in hospital environments (Harding *et al*, 2018). For *A. baylyi*, a non-pathogenic species belonging to the same genus, resilience against external stresses was shown to be at least partly attributed to extracellular polysaccharides (Ophir & Gutnick, 1994). These polysaccharides, whether secreted into the environment as biofilm matrix components (exopolysaccharides; EPS) or part of/attached to the bacterial membrane (like lipopolysaccharide [LPS], lipooligosaccharide [LOS], and capsule [capsular polysaccharide; CPS]), serve various protective roles against physical, chemical, and biological stresses (Flemming *et al*, 2023; Paczosa & Mecsas, 2016; Simpson & Trent, 2019; Whitfield *et al*, 2020). In *A. baumannii*, the capsule, encoded within the genomic region between the *fkpA* and *lldP* genes known as the K locus (Wyres *et al*, 2020), is a key feature for many strains. The K locus is usually arranged in three parts: I) the genes encoding the CPS export machinery (e.g., the *wza, wzb, wzc* operon); II) a central region for capsule construction and processing (including the genes *wzy* and *wzx,* which encode the repeat unit polymerase and translocase, respectively); and III) a module for synthesizing simple sugar substrates (Wyres *et al*., 2020). The CPS assembles into complex, multibranched glycans that are tightly anchored to the outer membrane by the Wzi protein, effectively encasing the cell in a protective polysaccharide shield (Tickner *et al*, 2021). This capsule plays a crucial role in the virulence of *A. baumannii*, as demonstrated in *in vivo* studies using animal models where it provided resistance against complement-mediated killing (Lees-Miller *et al*, 2013; Russo *et al*, 2010). Furthermore, monoclonal antibodies targeting CPS have been shown to protect mice from infection by hypervirulent strains (Nielsen *et al*, 2017). Despite its significant role in *A. baumannii* pathogenicity, the regulatory mechanisms governing capsule production remain largely unexplored. However, recent findings indicate that exposure to specific antibiotics at sub-MIC concentrations, such as of chloramphenicol, can trigger the upregulation of K locus genes (Geisinger & Isberg, 2015). This response is mediated by the BfmRS two-component regulatory system, leading to enhanced virulence of *A. baumannii* (Geisinger & Isberg, 2015).

Extracellular polysaccharides such as EPS and CPS are known to protect against attacks by bacteria with a type VI secretion system (T6SS), a key player in bacterial warfare (Smith *et al*, 2023). Found in 25 % of Gram-negative bacteria and in more than 50% of ß– and γ proteobacteria (Abby *et al*, 2016; Bingle *et al*, 2008), the T6SS is a contact-dependent contractile machinery that resembles inverted contractile bacteriophage tails (Basler *et al*, 2012; Leiman *et al*, 2009). The T6SS features a membrane complex that extends across both the inner and outer bacterial membranes, with a baseplate-like structure connected within the cytoplasm (Cherrak *et al*, 2018; Durand *et al*, 2015). The baseplate of the T6SS is linked to an internal tube composed of Hcp hexamers, encased by a contractile sheath formed by TssB and TssC proteins (Basler *et al*., 2012). Upon contraction, the T6SS propels its inner tube, along with VgrG-PAAR spike proteins and toxins, into neighboring cells, leading to either growth arrest or cell death (Cherrak *et al*, 2019; Russell *et al*, 2014). To protect themselves from the effects of their own T6SS-launched toxins, bacteria that possess the T6SS also produce immunity proteins. These proteins specifically neutralize the bacterium’s own toxic effector proteins, preventing self-intoxication or intoxication of kin (Hood *et al*, 2010; MacIntyre *et al*, 2010; Russell *et al*, 2011). As mentioned above, recent research has identified mechanisms of resistance to T6SS toxicity that don’t involve immunity proteins, which include defenses provided by the production of EPS (Granato *et al*, 2023; Hersch *et al*, 2020; Toska *et al*, 2018) and capsules (Flaugnatti *et al*, 2021).

The T6SS is widely found across *Acinetobacter* species, including *A. baumannii* (Dong *et al*, 2022; Weber *et al*, 2013). The genes encoding the core components of the T6SS reside in a single locus. However, the *vgrG* genes, which are crucial for the system’s function, are scattered throughout the chromosome alongside effector/immunity modules (Eijkelkamp *et al*, 2014; Lewis *et al*, 2019).

The regulation of the T6SS in *A. baumannii* varies, with some isolates expressing the system under standard laboratory conditions, while others regulate expression via proteins such as H-NS (Eijkelkamp *et al*, 2013; Repizo *et al*, 2015; Weber *et al*., 2013). Additionally, TetR-like repressors encoded on large conjugative plasmids, which also bear antibiotic resistance genes, can suppress T6SS to aid conjugation and plasmid dissemination among cells (Di Venanzio *et al*., 2019; Weber *et al*, 2015). This diversity in regulatory mechanisms indicates a complex interplay between antibiotic resistance, T6SS activity, and bacterial competitiveness. Indeed, the T6SS in *A. baumannii* significantly influences interbacterial dynamics, effectively targeting not only Gram-negative and Gram-positive bacteria (Le *et al*, 2021; Weber *et al*., 2013) but also exhibiting antifungal capabilities (Luo *et al*, 2023). However, although the T6SS confers competitive advantages to *A. baumannii* by targeting a wide range of microorganisms *in vitro*, studies in diverse animal models have shown that T6SS mutants do not incur a fitness cost (Weber *et al*., 2013), a finding that might be strain dependent, at least in the *Galleria mellonella* wax moth model of disease (Repizo *et al*., 2015). This suggests that the T6SS’s role in the bacterium’s virulence might not be direct (Subashchandrabose *et al*, 2015; Wang *et al*, 2014), highlighting a complex interaction with host organisms and/or its environment that warrants further investigation.

In this study, we investigate the impact of capsule production on T6SS antibacterial activity in *A. baumannii*. Our findings reveal that the capsular polysaccharide acts as a shield against T6SS attacks from rival bacteria. Despite this, many *A. baumannii* strains also have an operational T6SS, underscoring the capsules primary role as a one-way barrier. However, we show that under typical laboratory growth conditions, the presence of the surface-bound capsule nonetheless reduces the efficacy of the bacterium’s own T6SS. This T6SS impairment is further enhanced when CPS is overproduced due to genetic modifications or antibiotic treatment. Finally, we go on to demonstrate that when T6SS secretion is hindered, the accumulation of Hcp protein components is curtailed by a degradation process facilitated by the ClpXP protease system.

## Results and discussion

### The capsule of *A. baumannii* contributes to protection against T6SS attacks

In our study, we explored how capsule production influences the antibacterial activity of the T6SS in *A. baumannii*, specifically focusing on the clinical isolate A118 (Ramirez *et al*., 2010; Traglia *et al*, 2014). The A118 strain was found to possess the K locus, located between the *fkpA* and *lldP* genes (Vesel & Blokesch, 2021), suggesting its encapsulated nature. To confirm capsule production, we created a mutant lacking the *itrA* gene (Bai *et al*, 2021), essential for the initial steps of glycan chain formation (Kenyon & Hall, 2013). Analysis of the CPS material revealed the presence of high molecular weight polysaccharide in the wild-type (WT) strain but not the Δ*itrA* mutant, as indicated by Alcian blue staining (Fig. 1A). Next, we challenged both strains with rabbit serum to assess complement-mediated killing. As shown in Figure 1B, deleting the *itrA* gene resulted in a three-log decrease in survival compared to both the wild type (WT) strain and the mock control conditions. This complement-dependent killing of the non-capsulated strain was not observed when the serum was heat-inactivated before being added to the bacteria (Fig. 1B). Collectively, and in accordance with current literature (Kenyon & Hall, 2013; Lees-Miller *et al*., 2013), our findings establish the critical role of ItrA in CPS synthesis, and confirm that *A. baumannii* A118 possesses a capsule.

**Figure 1:**
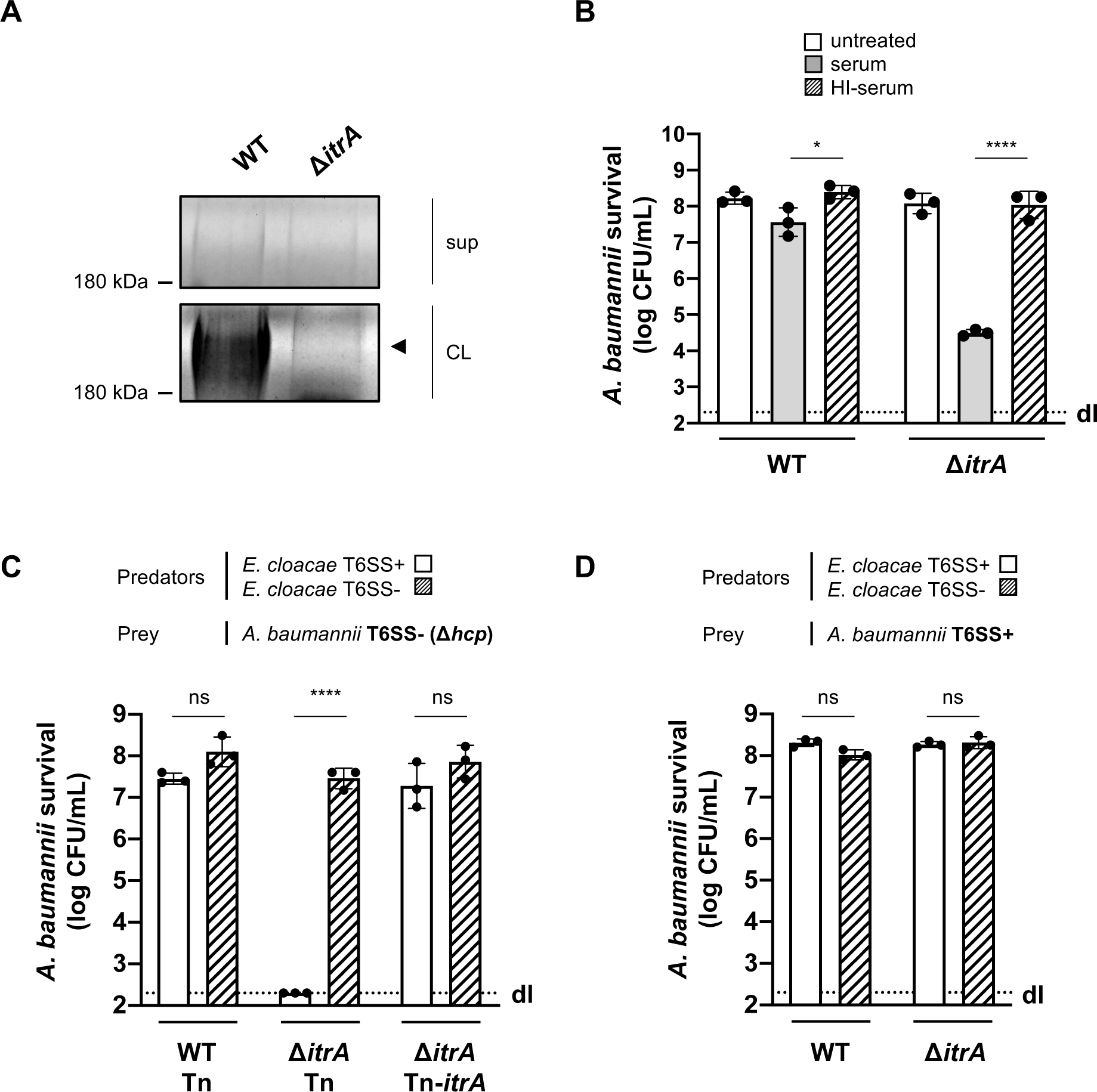
Capsular polysaccharide protects *A. baumannii* against external T6SS assaults. **(A)** Analysis of polysaccharides in the cell lysate (CL) and supernatant (sup) of wild-type (WT) or capsule-deficient (Δ*itrA*) strains of *A. baumannii,* separated by SDS-PAGE and stained with Alcian blue. The arrow indicates the polysaccharide band. **(B)** Protection against complement-mediated killing. Exponential growth cultures of WT and Δ*itrA* strains were incubated with PBS (untreated), complement-containing serum (serum), or heat inactivated serum (HI-serum) for 1 hour. Following treatment, the cultures were serially diluted and plated on LB agar to quantify colony-forming units (CFU), as shown on the *Y*-axis. **(C, D)** Capsule-dependent survival against T6SS assaults. T6SS-negative (Δ*hcp*) **(C)** or T6SS-positive **(D**) strains of *A. baumannii* were co-incubated with T6SS+ (white bars) or T6SS-(dashed bars) *Enterobacter cloacae*. Strains were either capsulated (WT background) or non-capsulated (Δ*itrA* background). In panel (C), capsulation was restored by provision of *P*_BAD_-*itrA* on a miniTn7 transposon (Tn-*itrA*) and provision of 2% arabinose. Tn is shown for WT and mutant strains containing the transposon without a specific cargo gene. *A. baumannii* survival was quantified and is shown on the *Y*-axis. The data represent means from three independent experiments with individual values shown by the circles (± SD, indicated by error bars). Statistical significance was assessed using an ordinary one-way ANOVA test. \**P* < 0.05, \*\*\*\**P* < 0.0001, ns = not significant. Detection limits (dl) were noted where applicable.

In our previous work, we observed that *A. baumannii* exhibited minimal susceptibility to *Enterobacter cloacae*’s T6SS (Flaugnatti *et al*., 2021), hinting at the potential protective role of its CPS against T6SS-mediated attacks. To investigate this protection further, we conducted a killing assay using capsulated (WT) and non-capsulated (Δ*itrA*) *A. baumannii* strains as prey in a T6SS-inactivated (T6SS-) strain background. Interestingly, the predator *E. cloacae* was ineffective in killing CPS-producing *A. baumannii* prey (WT carrying a cargo-less transposon; WT-Tn) in a T6SS-dependent manner (Fig. 1C). Conversely, the lack of CPS in the *A. baumannii* prey (Δ*itrA-*Tn) resulted in increased susceptibility to T6SS-mediated attacks by *E. cloacae* (Fig. 1C). This vulnerability could be reversed by introducing a new copy of the *itrA* gene into the strain’s genome (Δ*itrA*-Tn-*itrA*), as depicted in Figure 1C. When we conducted the experiment again under conditions that allow *A. baumannii*’s T6SS to function (T6SS+), we observed an unexpected outcome: the non-capsulated strain (Δ*itrA*) exhibited full resistance to *E. cloacae*’s T6SS attacks, mirroring the resistance shown by the capsulated WT strain (Fig. 1D). Indeed, several studies have reported that certain isolates of *A. baumannii* are equipped with a constitutively produced antibacterial T6SS (Repizo *et al*., 2015; Weber *et al*., 2013). To confirm the functionality of the T6SS in strain A118, we disrupted either the *hcp* or the *tssB* gene within the strain’s main T6SS gene cluster (Fig. S1A), which encode essential components of the system, and assessed the impact on T6SS-mediated antibacterial activity against *E. coli* prey (Fig. S1B). As expected, removing *hcp* and *tssB* effectively eliminated the T6SS’s antibacterial capabilities. These experiments verify that *A. baumannii* A118 indeed possesses an active antibacterial T6SS when tested under standard laboratory conditions. Collectively, our findings illustrate that both the capsule and the T6SS play pivotal roles in *A. baumannii*’s defense against T6SS-mediated attacks.

### CPS-deficient *A. baumannii* exhibits increased T6SS activity

The findings described above pose an intriguing question: How does *A. baumannii*’s CPS shield the bacterium from T6SS assaults by other microbes, yet still allow it to deploy its own T6SS weaponry? Or essentially, does the capsule function as a one-way barrier? To start addressing this question, we conducted a killing experiment with *A. baumannii* A118 strains, both CPS-positive (WT) and CPS-negative (Δ*itrA*), acting as predators. Initially, at a standard predator:prey ratio of 1:1, no significant differences in prey survival were observed between CPS-positive and CPS-negative strains, with survival rates at or below the detection limit (Fig. 2A). To enhance the sensitivity of the assay and increase its dynamic range, we adjusted the predator:prey ratio to 1:5, reflecting better invading bacteria. Under these conditions, we noted decreased T6SS-mediated killing activity in the WT strain compared to the Δ*itrA* strain (Fig. 2A). This suggests that T6SS activity is elevated in the absence of the CPS.

**Figure 2:**
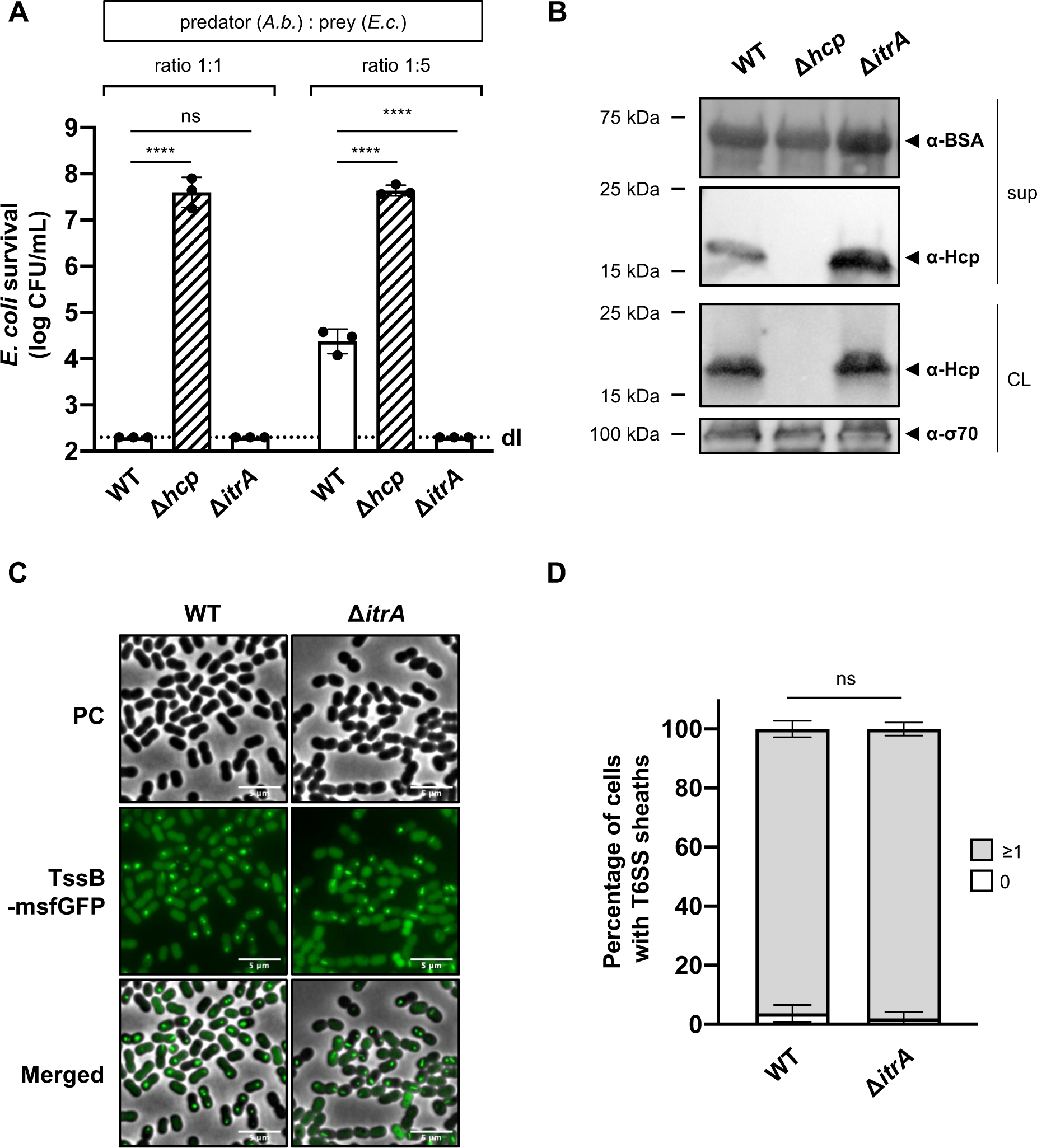
Capsular polysaccharide interferes with T6SS activity. **(A)** Non-capsulated strains show increased T6SS activity. Survival of *E. coli* (*E.c.*) after encountering capsulated (WT), T6SS-inactive (Δ*hcp*), or non-capsulated (Δ*itrA*) *A. baumannii* (*A.b.*), with different predator-to-prey ratios as indicated. Survival rates are shown as on the *Y*-axis. **(B)** Analysis of Hcp production and secretion in the strains mentioned in (A). Cell lysates (CL) and culture supernatants (sup) were tested through immunoblotting, using antibodies against Hcp (α-Hcp). The loading control (α-σ70) confirms equal amounts of the CL. BSA was added to supernatants and detected with α-BSA antibodies as a precipitation control. The data is representative of three independent experiments. **(C)** Fluorescence light micrographs of exponentially grown *A. baumannii* cells, either producing (WT) or not producing (Δ*itrA*) CPS, with a translational fusion (msfGFP) to the T6SS sheath protein TssB. Images include phase contrast (PC), green fluorescence (TssB-msfGFP), and a merged view of both channels. Scale bar: 5 μm. **(D)** Quantification of T6SS assembly over 5-minute time-lapses in TssB-msfGFP-carrying bacteria, comparing capsulated (WT; n=2832 cells) and non-capsulated (Δ*itrA;* n=2831 cells) cells. The *Y*-axis shows the percentage of cells producing T6SS structures, with cells not producing T6SS in white and those producing at least one structure in gray. Data are averages from three experiments (± SD, as defined by error bars). Statistical significance compared to WT is marked, determined via an ordinary one-way ANOVA test **(A)** or a two-way ANOVA test **(D)**, with \*\*\*\**P* < 0.0001, ns = not significant. Detection limits (dl) are indicated.

To verify the increased T6SS activity in the absence of CPS, we performed an Hcp secretion assay. This assay, a benchmark for assessing T6SS functionality (Pukatzki *et al*, 2006), relies on immunodetecting the Hcp protein in the supernatant, with intracellular Hcp serving as a control. Importantly, while both strains produced Hcp at comparable levels, the amount of Hcp detected in the supernatant was considerably higher in the Δ*itrA* strain (Fig. 2B). This finding reinforces the notion that T6SS activity is enhanced in the absence of the capsular polysaccharide. As an internal BSA precipitation control ensured that the observed differences in Hcp recovery were not due to variations in precipitation efficiency, we hypothesized that the CPS might directly impact the assembly of and/or firing by the T6SS machinery. We therefore compared T6SS structures within cells using a functional translational fusion between TssB and msfGFP (Fig. S1B), as previously reported (Lin *et al*, 2022). To objectively assess T6SS assembly, we developed a tool designed for the automatic analysis T6SS structures in cells over a 5-minute time interval (Movie S1). Our observations revealed highly dynamic T6SS structures in nearly all WT (96.2 % ± 2.8) and *itrA* mutant cells (98.0% ± 2.2) (Figs. 2C and D). This data indicates that the capsule’s presence or absence does not affect the production or assembly of the T6SS in *A. baumannii* A118.

Collectively, our findings indicate that CPS does not hinder the secretion process of the T6SS or the consequent elimination of competing cells. However, we also uncovered that the capsule modulates T6SS activity, as shown by the variations in killing efficiency and Hcp secretion between encapsulated and non-encapsulated strains. This suggests that the capsule may serve as an additional barrier the T6SS has to traverse to be expelled from the cell. Supporting this theory, our analysis reveals that the enhanced T6SS activity in the non-capsulated mutant (Δ*itrA*) is not due to a higher number of T6SS assemblies but likely due to an increase in the number of successful T6SS firing events. This finding is in line with previous reports on *Campylobacter jejuni*, where the T6SS was only cytotoxic to red blood cells in a capsule-deficient context (Bleumink-Pluym *et al*, 2013), leading to the hypothesis that the capsule acts as a physical barrier, limiting T6SS’s ability to directly interact with target cells. Variations in capsule production have been observed in *A. baumannii*, which employs a kind of bet-hedging strategy that leads to the formation of two types of variants within the same clonal population, namely opaque and translucent colonies. These variants are capable of phenotypically switching between these states, thereby enhancing their adaptation to diverse environments (Chin *et al*, 2018). Such a strategy in capsule modulation can offer significant advantages, including protection against external threats like complement-mediated killing, as well as competitive interactions with surrounding organisms. Furthermore, the observed increase in T6SS activity in the non-capsulated strain suggests a compensatory mechanism for the absence of the protective capsule layer.

### Alterations in the organization of capsule material disrupt the secretion process

Having established that CPS interferes with the T6SS secretion process, we next explored whether enhancing CPS production could entirely block T6SS activity. Previous research has identified two genetic alterations that increase CPS secretion and/or production (Geisinger & Isberg, 2015). Indeed, Geisinger and Isberg demonstrated that a substitution in the Walker A motif of the Wzc protein, which controls the size of exported polysaccharides, induces in a mucoviscous phenotype characterized by abnormally high molecular weight polysaccharides predominantly found in the supernatant and only loosely attached to the cell. The second mechanism for elevated CPS production involves the two-component system BfmRS, recognized for its role in various cellular processes including biofilm formation, serum resistance, antibiotic resistance, and envelope stress response (Geisinger *et al*, 2018; Russo *et al*, 2016; Tomaras *et al*, 2008). The BfmS histidine kinase within this system typically represses K locus expression by phosphorylation of the response regulator BfmR (Palethorpe *et al*, 2022). Consequently, removing *bfmS* disrupts this phosphorylation cascade, resulting in the overproduction of the K locus gene cluster (Geisinger & Isberg, 2015).

To enhance CPS production in *A. baumannii* A118, we therefore introduced a point mutation in *wzc* encoding the Walker A motif variant [K547Q], and we also created a deletion mutant of *bfmS*. Both modifications led to the formation of mucoviscous colonies (Fig. S2A-B), with a noticeable difference: stretching of the Wzc[K547Q] colonies produced a string (> 5mm), a phenomenon not observed in the mucoviscous Δ*bfmS* mutant. This suggests that the capsular characteristics differ between the two mutants. Indeed, further analysis, including CPS extraction followed by Alcian blue staining and the serum-mediated killing assay, revealed distinct outcomes for these mutants. The Δ*bfmS* mutant showed increased resistance to serum-mediated killing (Fig. 3A), aligned with an augmented presence of cell-associated CPS material (Fig. 3B). Conversely, the Wzc[K547Q] mutant displayed heightened susceptibility to the rabbit serum (Fig. 3A), which correlates with the faint CPS signal detected in the cell fraction by Alcian blue staining (Fig. 3B). This indicates that dysregulation in polysaccharide chain length can adversely affect the capsule’s protective properties.

**Figure 3:**
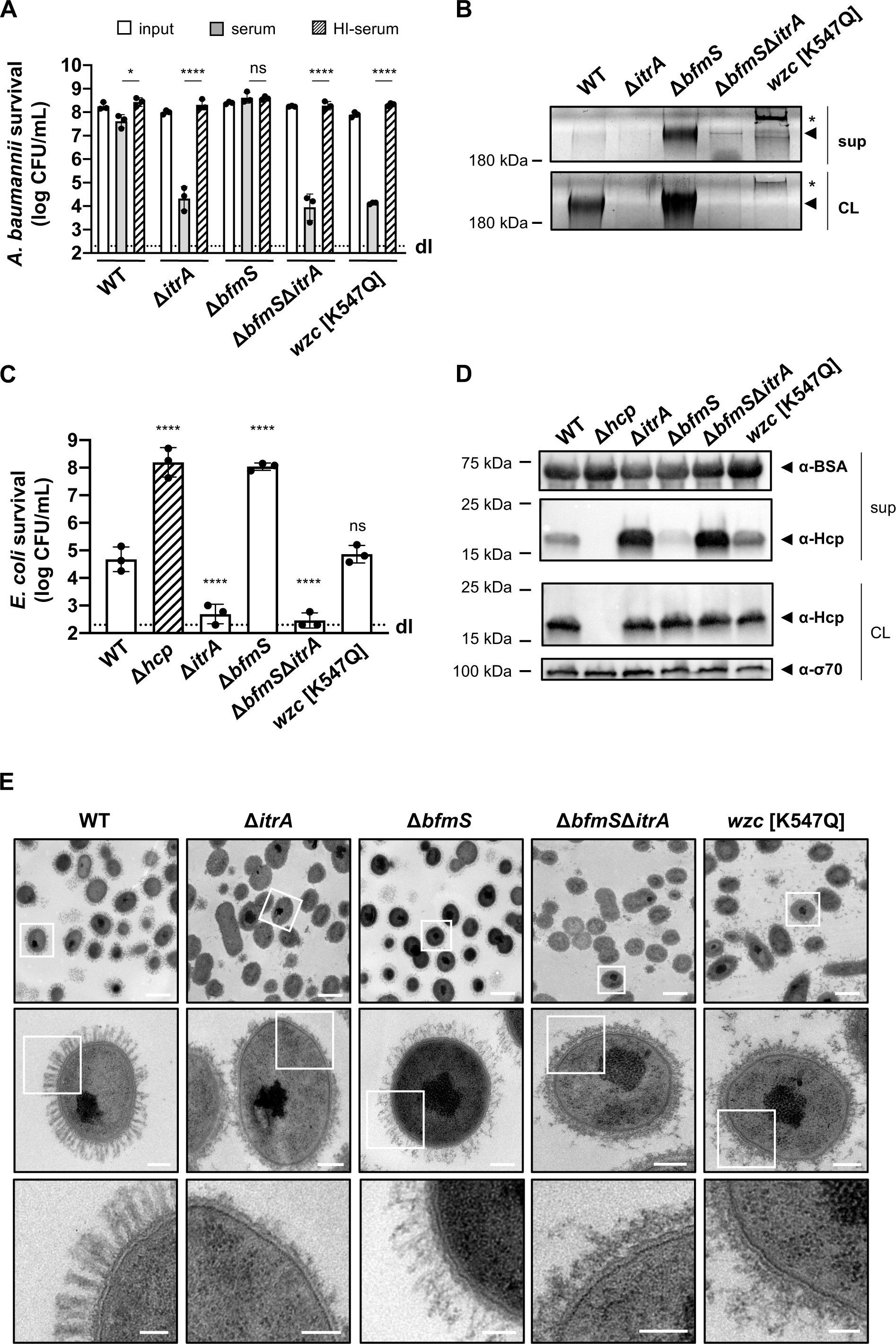
Increased CPS production inhibits T6SS activity. **(A)** Complement resistance assay across *A. baumannii* strains. The assay tested the resistance against complement-containing serum of these strains: capsulated wild-type (WT), non-capsulated (Δ*itrA*), capsule-overproducing (Δ*bfmS*), a Δ*bfmS*Δ*itrA* double mutant, and a strain carrying mutated *wzc* (encoding Wzc[K547Q]). Details as described for Figure 1B. **(B)** Polysaccharide analysis in wild-type (WT) and variants described in panel A. Polysaccharides from cell lysate (CL) or supernatants (sup) were separated by SDS-PAGE and stained with Alcian blue. Arrows point to polysaccharide bands with the asterisks marking high molecular size polysaccharides. **(C)** Survival of *E. coli* prey after interaction with the *A. baumannii* strains described in panel (A) as predators. A T6SS-inactive strain (Δ*hcp*, dashed bar) was added as control. The predator-to-prey ratio of 1:5 was used. Survival rates are indicated on the *Y*-axis. Details as for Figure 2A. **(D)** Hcp production and secretion levels of WT and mutant *A. baumannii* strains described in panel **(A).** Details as in Fig. 2B. **(E)** Transmission Electron Microscopy images of WT, Δ*itrA*, Δ*bfmS*, Δ*bfmS*Δ*itrA,* and *wzc*[K547Q] strains. White squares indicate zoomed areas. Scale bars correspond to 1 μm, 0.2 μm, and 0.1 μm for the top, middle, and bottom images, respectively. Data for panels (**B)**, (**D**), and (**E**) are representative of three independent experiments. For panels (**A**) and (**C**), data points are averages from three experiments (± SD, shown by error bars). Statistical significance compared to the heat-inactive serum treatment (**A**) or to the WT strain (**C**) is noted above the charts, determined with an ordinary one-way ANOVA test. \**P* < 0.05, **** *P* < 0.0001, ns = not significant. Detection limits (dl) were noted where applicable.

To assess the impact of these altered CPS profiles on T6SS activity, we utilized these mutants as predators in a killing assay. The results reveal that the Δ*bfmS* mutant exhibits a significant reduction in its ability to kill *E. coli* prey cells (Fig. 3C). By complementing the Δ*bfmS* mutant *in cis* with a copy of *bfmS* under its native promoter, T6SS-mediated killing was restored to levels similar to those of the WT strain (Fig. S2C). To rule out the possibility of the *bfmS* mutation having broad effects on T6SS production or function, we also evaluated the double mutant Δ*bfmS*Δ*itrA*, which reinstated the strain’s killing ability (Fig. 3C). These findings were consistent with the Hcp secretion profiles of these mutants. Specifically, the mutant lacking *bfmS* showed a significant impairment in its Hcp secretion activity, whereas the Δ*bfmS*Δ*itrA* double mutant reflected the secretion pattern of the Δ*itrA* strain (Fig. 3D). This suggests that the observed phenotype indeed results from the increased production of the capsule.

To investigate whether capsule-secreted material could interact with proteins in the supernatant, we co-cultured the WT and a secretion-impaired Δ*tssB* mutant and compared their Hcp secretion profiles to the WT co-cultured with Δ*bfmS.* We observed no significant differences in the levels of Hcp protein secretion between the two conditions (Fig. S2D), indicating that the secretion defect seen in Δ*bfmS* is attributable to an impairment in secretion rather than to interactions with CPS in the supernatant. We also engineered a *bfmS* deletion in two environmental *A. baumannii* isolates (29D2 and 86II/2C) (Wilharm *et al*, 2017). As illustrated in Figure S2E, both strains are capable of producing an antibacterial T6SS. Importantly, the deletion of *bfmS* in these strains also resulted in the inhibition of the T6SS-mediated killing. These findings indicate that *bfmS* affects T6SS activity across different strains.

Unlike the Δ*bfmS* phenotypes, the Wzc[K547Q] variant demonstrated T6SS-mediated killing similar to that observed in the wild type (WT), as shown in Figure 3C. Consistently, this variant also exhibited Hcp secretion levels that appeared comparable (or even increased) to those of the WT (Fig. 3D). This finding was unexpected, as an increase in CPS length might be presumed to hinder T6SS activity. However, this outcome is in line with the observation that the Wzc[K547Q] variant was not protected from complement-mediated killing (Fig. 3A).

To gain more insight into the ultrastructure of the capsule in the different genetic backgrounds, we imaged cells using transmission electron microscopy (TEM) (Fig. 3E). The WT cells were surrounded by material forming large finger-like projections extending about 150 nm from the cell surface, arranged in a semi-regular pattern of projections and spaces. As expected, these structures were absent in the Δ*itrA* mutant, confirming its essential role in capsule assembly. Notably, the Wzc[K547Q] variant also lacked these structures, appearing similar to the Δ*itrA* mutant. However, we observed a significant presence of what is presumed to be capsular material floating in the medium surrounding the cells, with additional material potentially being lost during the fixation process (Fig. 3E). This detached CPS aligns with the Alcian blue staining results (Fig. 3B) and could explain the observed differences in the string test results for the Wzc[K547Q] colonies compared to the Δ*bfmS* mutant (Fig. S2A). The creation of such a viscous environment by the release of long-chain CPS may therefore impact T6SS activity, explaining the decreased killing ability compared to the Δ*itrA* mutant (Fig. 3C). In contrast, the *bfmS* mutant exhibited a dense, tangled, mesh-like network of CPS covering the cell surface, similar to the wild type (WT) but without the clear periodic spaces. As expected, this dense capsule network was absent in the Δ*bfmS*Δ*itrA* double deletion mutant (Fig. 3E).

Taken all together, these results indicate that disruption in the organized, finger-like structure of the capsule, as seen with overexpression of the K-locus, leads to a suppression of T6SS activity and blocks Hcp secretion. This observation highlights the importance of CPS’s surface organization in affecting the extracellular secretion process. It is tempting to speculate that, within the WT scenario, T6SS may deploy through gaps akin to arrow-slit in the capsule’s mesh, a process that becomes unfeasible when CPS organization is disrupted. This concept mirrors a hypothesis suggested by Toska and colleagues for *Vibrio cholerae,* where T6SS secretes through biofilm-associated exopolysaccharide (Toska *et al*., 2018). An alternative explanation might be that capsule overexpression enhances polysaccharide dispersion into the surroundings. Coupled with changes to the capsule directly attached to the cell surface, this could effectively increase the spatial gap between cells, impeding T6SS functionality.

### Antibiotics-induced CPS production impairs T6SS activity

It has been shown that the *A. baumannii* isolate 17978 boosts CPS production via the BfmRS two component system in response to sub-minimal inhibitory concentrations (sub-MIC) of chloramphenicol (Geisinger & Isberg, 2015). When we exposed *A. baumannii* A118 to various chloramphenicol concentrations, we found that the capsule induction by the antibiotic was dose-dependent, as evidenced by increased CPS presence in the supernatant (Fig. 4A). We next asked whether T6SS activity inhibition seen in the Δ*bfmS* mutant could also be induced under antibiotic-triggered capsule overproduction conditions in the WT background. Unfortunately, the Hcp secretion assay did not yield conclusive results due to contamination from cytoplasmic material, indicating that chloramphenicol exposure led to partial cell lysis (Fig. 4B). However, we noticed enhanced T6SS-mediated killing in the non-capsulated strain (Δ*itrA*) versus the capsulated (WT) under antibiotic exposure (Fig. 4C). This suggests that chloramphenicol-induced capsule production disrupts T6SS activity. It is important to note that antibiotic treatment alters *A. baumannii*’s growth, which may change the predator:prey ratio during the assay and affect the experimental results. However, this factor should similarly impact the Δ*itrA* mutant, which demonstrates effective killing even in the presence of chloramphenicol.

**Figure 4:**
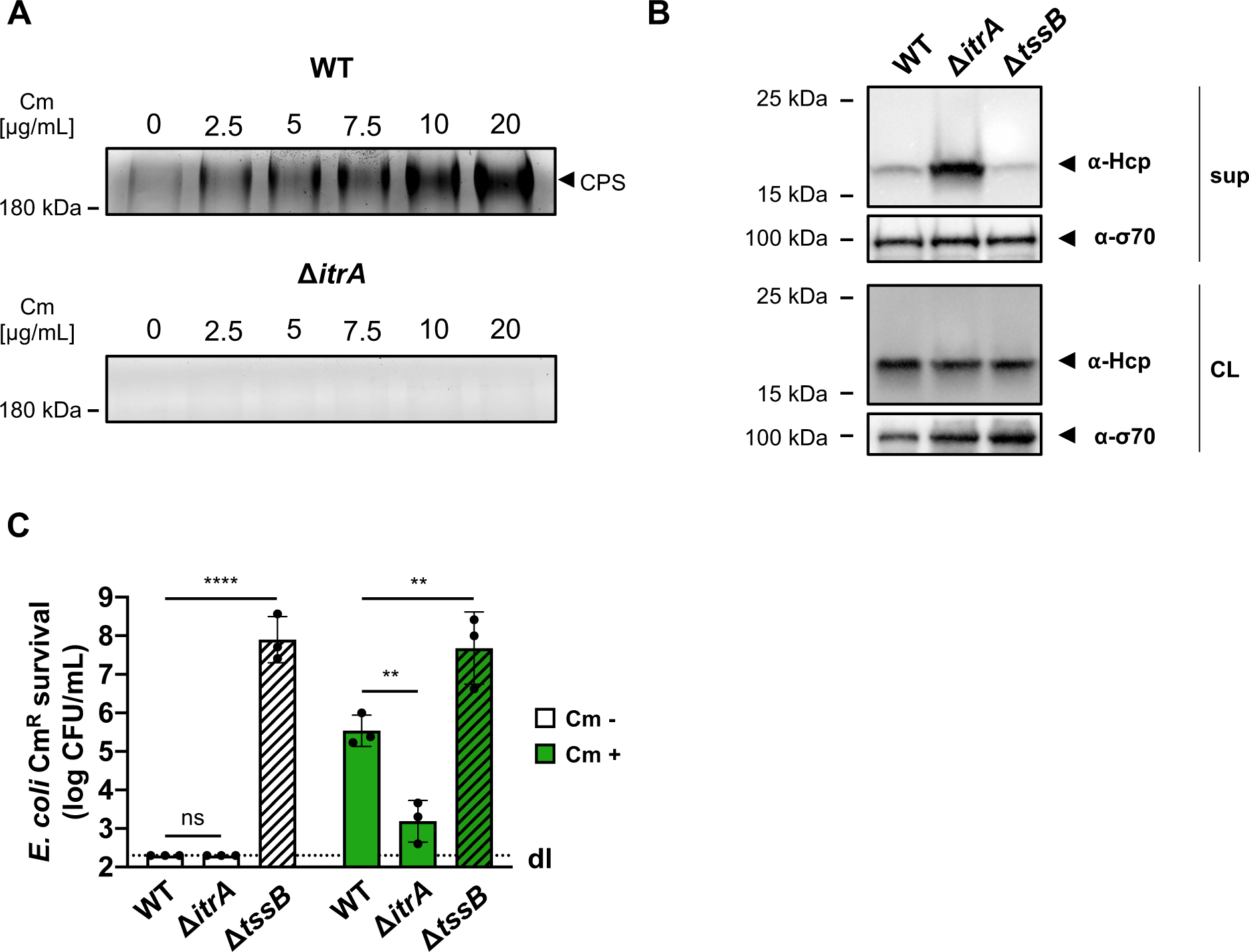
T6SS inhibition upon CPS overproduction due to antibiotic treatment. **(A)** Capsule production in wild-type (WT, upper panel) or capsule-deficient (Δ*itrA*, lower panel) strains was induced using varying concentrations of chloramphenicol, as indicated. Polysaccharides from supernatant were precipitated, separated by SDS-PAGE, and visualized with Alcian blue staining. The arrow marks the polysaccharide band. (**B)** Hcp secretion in chloramphenicol-treated cells. Hcp production and secretion were analyzed in WT, Δ*itrA*, and Δ*tssB* strains through immunoblotting. Details as described in Figure 2B. σ-70 detection served as loading (CL) and lysis control (sup). **(C)** Survival of chloramphenicol-resistant (Cm^R^) *E. coli* after contact with capsulated (WT), non-capsulated (Δ*itrA*), or T6SS-inactive (Δ*tssB*, dashed bars) *A. baumannii* predators, either unexposed (white bars) or exposed to 20μg/mL chloramphenicol (green bars) to induce capsule production. Survival rates are indicated on the *Y*-axis. For panel (**C**), data points are averages from three independent experiments (± SD, shown by error bars). Statistical significance was determined with an ordinary one-way ANOVA test. \*\**P* < 0.01, \*\*\*\**P* < 0.0001, ns = not significant. Detection limits (dl) were noted where applicable.

Collectively, this data suggests that the inhibition of T6SS by increased capsule production, as observed with the Δ*bfmS* mutant, could be relevant in natural conditions that *A. baumannii* might encounter, such as the presence of sub-MIC antibiotics in the environment. Indeed, this mucoid state has been observed with other antibiotics apart from chloramphenicol, some of which are used in clinical settings (Geisinger & Isberg, 2015; Traub & Bauer, 2000). Interestingly, Geisinger and Isberg demonstrated that the antibiotic-induced enhancement of capsule production represents a non-mutational phenotype, which can be reversed upon removal of the antibiotic (Geisinger & Isberg, 2015). It is therefore tempting to speculate that this inverse relationship between capsule production and T6SS activity may provide adaptive advantages in response to environmental changes and competitive interactions with other bacteria, as proposed by Weber and colleagues (Weber *et al*., 2015).

### Sensory function is preserved in capsule-overproducing strains

A recent investigation into *Acinetobacter baylyi* revealed the presence of TslA, a periplasmic protein essential for precise assembly of the T6SS machinery at points of contact with other cells, aiming to prevent wasteful T6SS firing events (Lin *et al*., 2022). Importantly, the periplasmic *Acinetobacter* type six secretion system-associated A protein (AsaA), which is the TslA homolog in *A. baumannii*, has been shown to play a role in efficient T6SS activity (Li *et al*, 2019). Indeed, Li *et al*. suggested that AsaA/TslA impacts the assembly or stability of the T6SS through its interaction with the membrane complex protein TssM (Li *et al*., 2019). These findings led to the speculation that in the Δ*bfmS* mutant, the elevated levels of capsule production could obstruct the environmental sensing function of TslA, thereby reducing the likelihood of T6SS assembly.

To test this hypothesis, we analyzed the dynamics of the TssB-msfGFP fusion in WT, Δ*tslA,* and Δ*bfmS* backgrounds using time-lapse microscopy over a five-minute time span (Fig 5A-B). We noted a significant 5.6-fold decrease in T6SS assembly and activity in the Δ*tslA* mutant (17.6 ± 8.7 %) compared to the WT (99.0 ± 1.4 %). Meanwhile, the Δ*bfmS* mutant exhibited a more moderate 1.9-fold reduction (50.8 ± 8.8 %) in comparison to the WT. Remarkably, the Δ*bfmS*Δ*itrA* double mutant showed T6SS assembly rates (97.6 ± 1.0 %) similar to both the WT and Δ*itrA* (99.2 ± 1.0 %) strains. These findings suggest that the capsule’s overproduction in Δ*bfmS* only partially influences T6SS assembly. Of note, the observed decrease in fluorescence intensity in the Δ*bfmS* and Δ*bfmS*Δ*itrA* strains remains unexplained. Nonetheless, analysis confirmed that the fusion protein levels were consistent across all strains (Fig. S2F), indicating that TssB-msfGFP production is unaffected in these mutants.

**Figure 5:**
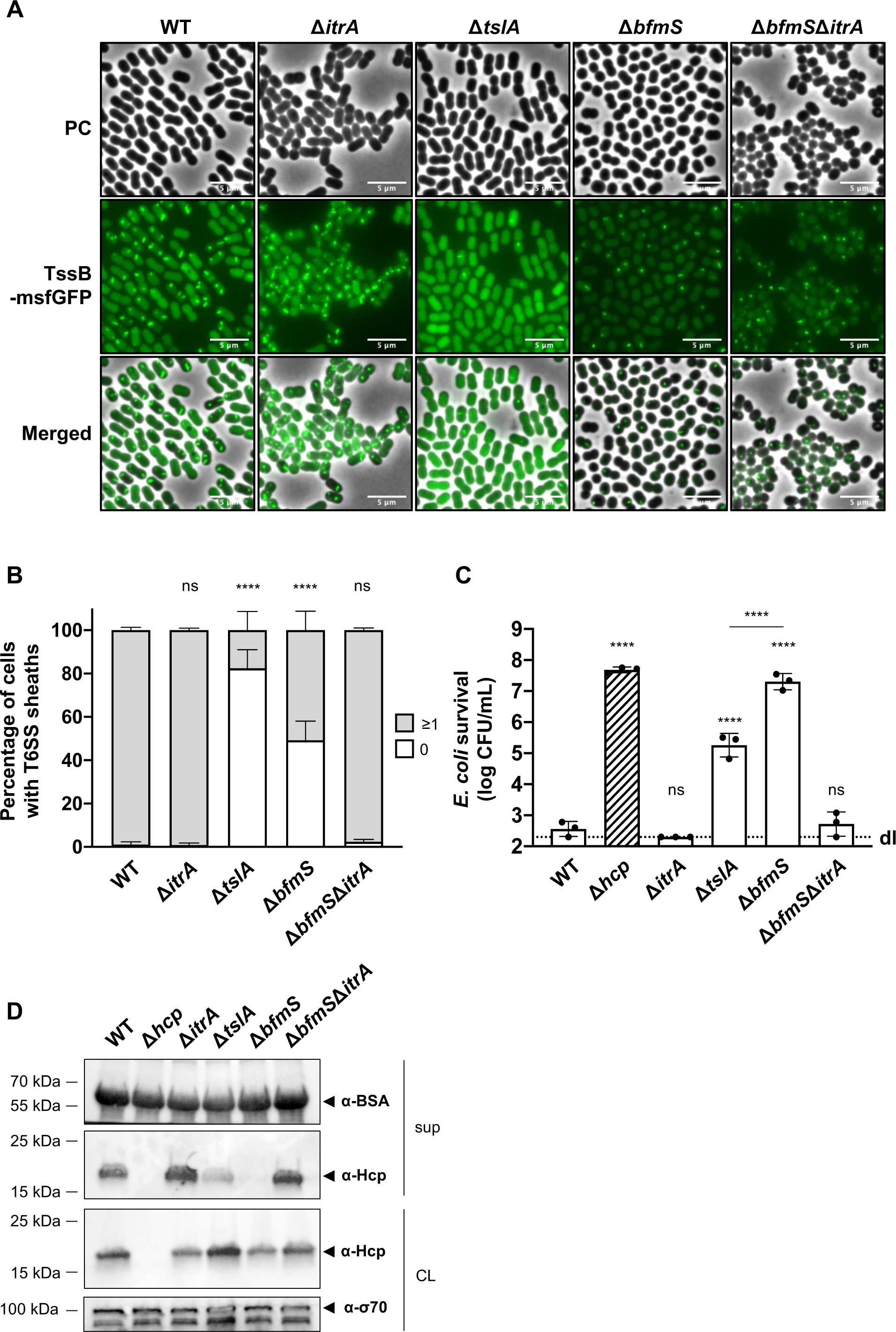
T6SS inhibition in CPS overproducing strain goes beyond inability of cell-to-cell contact sensing. **(A)** Fluorescence light micrographs of TssB-msfGFP-producing *A. baumannii*. Strain backgrounds: capsulated (WT), non-capsulated (Δ*itrA*), cell contact sensing mutant (Δ*tslA*), capsule overexpressing (Δ*bfmS*), and Δ*bfmS*Δ*itrA* double mutant. Details as described for Figure 2C. Scale bar: 5 μm. **(B)** Quantification of T6SS structures in the *A. baumannii* strains described in panel (A). Details as for Figure 2D. Number of analyzed cells was 3041, 2685, 2805, 3667, and 4800 for the strains indicated on the *X*-axis. Data are averages from three independent experiments (± SD, as defined by error bars). (**C**) Survival rates of *E. coli* prey after exposure to the *A. baumannii* WT, Δ*itrA*, Δ*tslA*, Δ*bfmS,* and Δ*bfmS* predator strains with native (non-fused) *tssB.* A predator-to-prey ratio of 1:1 was used. Survival is indicated on the *Y*-axis. Bars indicate mean values (± SD, as shown by error bars). **(D)** Hcp production and secretion were analyzed for the same *A. baumannii* strains as in panel (**C**). Experimental details as for Figure 2B. Statistical analyses show the significance compared to WT conditions, utilizing a two-way ANOVA test for (**B**) and an ordinary one-way ANOVA test for (**C**). \*\*\*\**P* < 0.0001, ns = not significant. Detection limits (dl) are indicated.

To further investigate how the absence of contact sensing affects T6SS secretion activity, we delved into the role of TslA in *A. baumannii* A118 by conducting a killing experiment (Fig. 5C) and a Hcp secretion assay (Fig. 5D). In line with results obtained previously in *A. baylyi* (Ringel *et al*, 2017) and *A. baumannii* ATCC17978 (Kandolo *et al*, 2023; Li *et al*., 2019), the removal of *tslA* led to a decrease in T6SS-mediated killing and the amount of secreted Hcp. Interestingly, the *tslA* mutant displayed significantly higher levels of killing (Fig. 5C) and Hcp secretion (Fig. 5D) compared to the Δ*bfmS* mutant under identical experimental conditions.

Taken together, these findings collectively indicate that the diminished killing and secretion performance seen in the Δ*bfmS* mutant cannot be solely attributed to a defect in cell-cell contact sensing. Despite assembling around three times more T6SS structures compared to the Δ*tslA* mutant, the Δ*bfmS* mutant exhibits a T6SS killing activity that is 100 times less effective. This evidence points to the conclusion that T6SS assembly is not the limiting factor for the T6SS inhibition in the Δ*bfmS* mutant.

### Prolonged secretion inhibition triggers Hcp degradation

Given that our data indicate an inhibition of T6SS activity by CPS upregulation, we explored the possibility of regulatory cross-talk between these two processes during prolonged capsule overproduction. To address this question, we assessed Hcp production levels in various capsule mutant backgrounds using strains in stationary phase (15-16 h of growth) (Fig. 6A). Our observations revealed that, in stationary phase cultures, Hcp was undetectable in the Δ*bfmS* mutant but present at WT levels in the Δ*bfmS*Δ*itrA* double mutant (Fig. 6A). This contrasts with results from exponentially growing cultures, where all strains produced Hcp at comparable levels (Fig. 3D). Given the impaired Hcp secretion in the Δ*bfmS* mutant, we hypothesized that intracellular accumulation of Hcp or another unidentified signal may trigger a feedback mechanism that down-regulates Hcp production.

**Figure 6:**
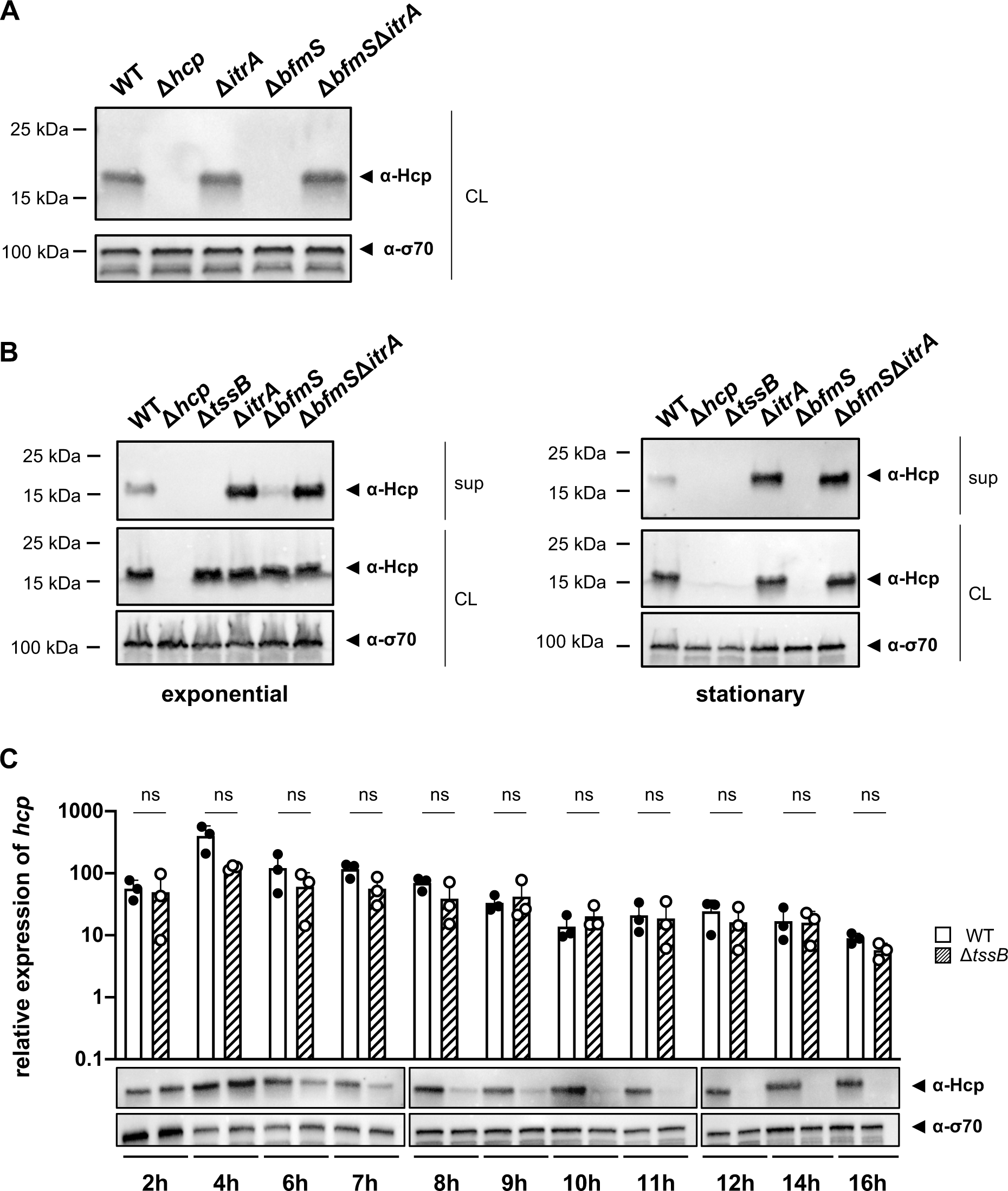
Secretion-impaired strains degrade Hcp during stationary phase. **(A)** Immunoblot analysis of Hcp protein levels in cell lysates (CL) of capsulated (WT), T6SS-inactive (Δ*hcp*), non-capsulated (Δ*itrA*), capsule overexpressing (Δ*bfmS*) and Δ*bfmS*Δ*itrA* mutant strains grown to stationary phase. (**B**) Comparative analysis of Hcp production and secretion in *A. baumannii* strains during exponential (left) and stationary (right) growth phases. Strains as explained in panel **(A)** with the addition of a secretion-impaired Δ*tssB* mutant. Details as described for Figure 2B. **(C)** Hcp abundance is regulated at the post-translational level. The graph shows relative *hcp* gene expression levels over a 16-hour period in the WT (white bars) versus the secretion-impaired strain (Δ*tssB*, dashed bars). The lower panel illustrates Hcp protein production over the same time frame, analyzed by immunoblotting. These results are representative of three independent experiments, and the bars show the mean (± SD, as defined by error bars). Statistical analyses were performed on log-transformed data using a two-way ANOVA. ns = not significant.

Interestingly, a recent study reported that *V. cholerae* can sense Hcp levels and regulate T6SS expression accordingly (Manera *et al*, 2021). The authors demonstrated that the RpoN-dependent regulator VasH interacts with Hcp, influencing the expression of auxiliary T6SS clusters that include the *hcp* genes. However, unlike *V. cholerae*, the main T6SS cluster in *A. baumannii* does not contain genes for bacterial enhancer binding proteins (bEBP) such as VasH (Fig. S1A), indicating a potentially different regulatory mechanism. To further explore the effects of Hcp accumulation in *A. baumannii*, we employed the secretion-deficient Δ*tssB* mutant. We measured Hcp levels under two different growth conditions: exponential and stationary phases (Fig. 6B). Surprisingly, similar to the Δ*bfmS* mutant, deletion of *tssB* led to the disappearance of Hcp in stationary phase. Comparable results were obtained with secretion-impaired mutants of *A. baumannii* strains 29D2 and 86II/2C (Fig. S3A), suggesting that Hcp downregulation upon blocking secretion might be a common feature in *A. baumannii*. These findings suggest that the reduced T6SS assembly and activity observed in the Δ*bfmS* mutant increase the cytoplasmic pool of Hcp protein, which then triggers the down-regulation or degradation of Hcp.

To determine if the observed phenotype was specific to Hcp or affected other T6SS components, we assessed TssB-GFP levels using GFP antibodies in the Δ*bfmS* strain (Fig. S3B). We observed no significant reduction in TssB-GFP levels in the Δ*bfmS* strain compared to other strains carrying *tssB-msfgfp*, suggesting that the downregulation or degradation might be specific to Hcp. To further understand the timing of this phenotype, we monitored Hcp protein production and *hcp* mRNA levels in both the WT and the secretion-impaired Δ*tssB* mutant over a 16-hour period (Fig. 6C). Notably, the decrease in Hcp protein levels in Δ*tssB* occurred between 6 and 11 h of growth, coinciding with the transition to late stationary phase (Fig. S3C). However, there were no statistically significant changes in *hcp* transcript levels between the WT and the Δ*tssB* mutant throughout the experiment. These results suggest that the effects of secretion impairment and Hcp intracellular accumulation may be regulated post-transcriptionally.

To further explore this regulatory mechanism, we attempted to overexpress Hcp in both the WT and Δ*hcp* backgrounds, monitoring *hcp* mRNA levels and Hcp protein production. Upon induction, *hcp* mRNA levels increased significantly, leading to elevated Hcp protein levels in the WT background (WT + p-*hcp*) (Fig. 7A). However, in the Δ*hcp* background (Δ*hcp* + p-*hcp*), Hcp protein production was not detectable under these conditions despite high transcript levels (Fig. 7A). When *hcp* was overexpressed in *E. coli* as a control condition, both the transcript and the protein were successfully produced and detected (Fig. S3D), similar to the situation in the WT *A. baumannii* background.

**Figure 7:**
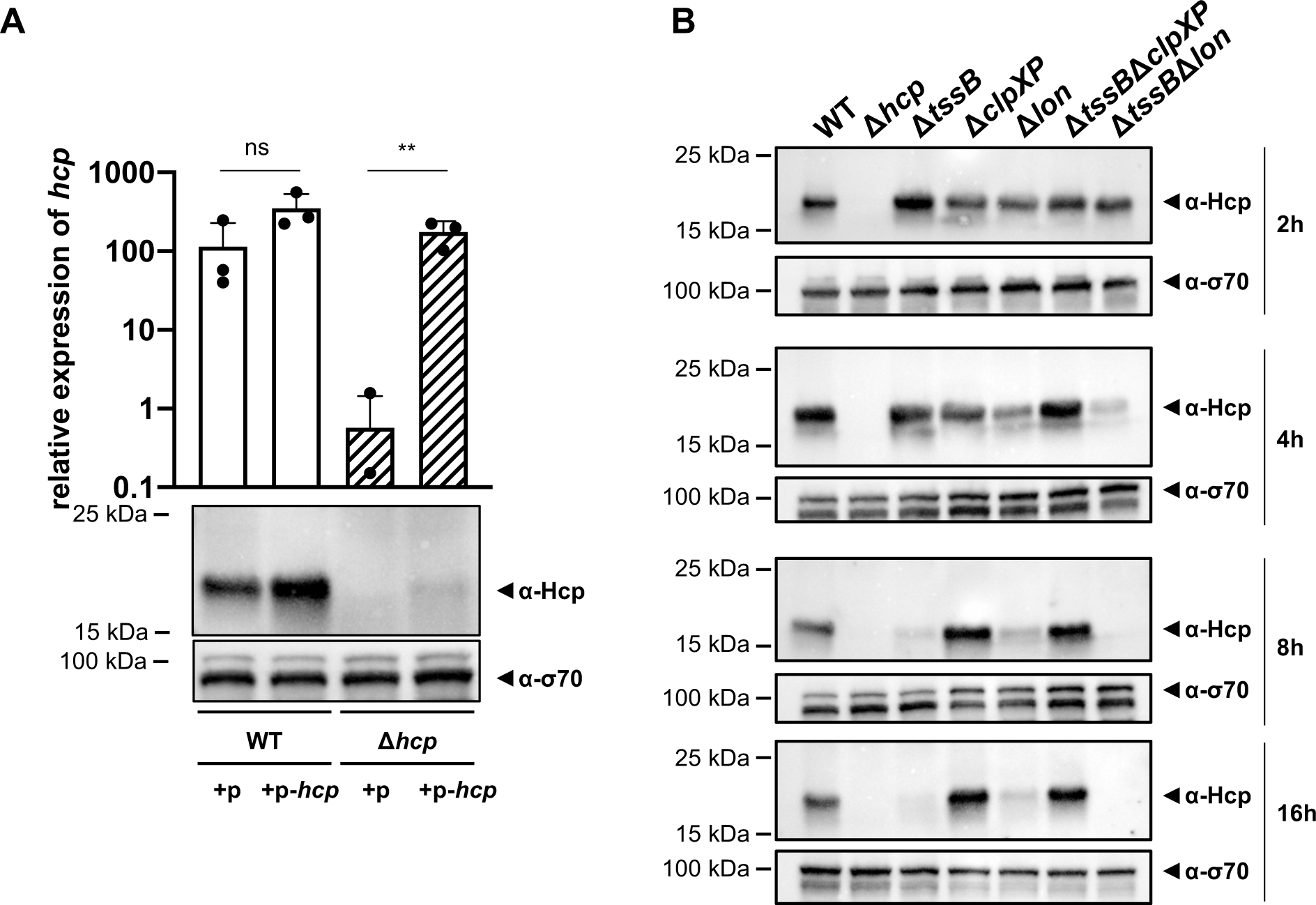
ClpXP protease machinery mediates Hcp degradation. **(A)** Hcp levels are low in secretion-impaired strains. The upper panel shows the relative expression levels of *hcp* in WT or the *hcp* mutant, either carrying an empty plasmid (+p) or a plasmid designed for *hcp* overexpression (+p-*hcp*). The panel below the graph illustrates Hcp abundance in these strains, as assessed by immunoblotting. **(B)** Hcp accumulation over time in *clpXP*-deficient mutants. Hcp levels were examined in cell lysates from various *A. baumannii* strains including capsulated WT, T6SS-inactive (Δ*hcp* and Δ*tssB*), and protease-deficient (Δ*clpXP* and Δ*lon*) mutants, or strains lacking multiple genes (Δ*tssB*Δ*clpXP* and Δ*tssB*Δ*lon*). The bacteria were grown over 2, 4, 8, and 16-hour growth periods. Immunoblot analyses were performed as described for Figure 2B. Data from three independent experiments are presented as means (± SD, as shown by error bars). Statistical significance was assessed on log-transformed data using a two-way ANOVA. \*\**P* < 0.01, ns = not significant.

The findings suggest that Hcp is regulated at the post-transcriptional level, potentially through a degradation mechanism. ClpXP and Lon proteases are known to play crucial roles in various stress responses, specifically in degrading misfolded or accumulated proteins to mitigate proteotoxic stress (Sauer & Baker, 2011). To investigate the potential involvement of these general proteases in the post-transcriptional regulation of Hcp, we generated individual Δ*clpXP* and Δ*lon* mutants, as well as combinations with a secretion-impaired background (Δ*tssB*Δ*clpXP* and Δ*tssB*Δ*lon*). Intriguingly, while the deletion of *lon* had no discernible effect, we detected Hcp protein in the Δ*tssB*Δ*clpXP* double mutant even at late stages of growth, indicating that ClpXP may be involved in the degradation mechanism of Hcp observed when secretion is impaired (Fig. 7B).

Given the time-dependent nature of this degradation, we analyzed the transcript levels of *clpX* and *clpP* at various times (2h, 6h, 8h, and 12h) before, during, and after the degradation of Hcp, both under secretion-permissive (WT) and non-permissive (Δ*tssB*) conditions (Fig. S3E). We observed an increase in *clpX* transcripts at 6h, corresponding with the transition into late stationary phase (Fig. S3C). However, there were no statistically significant differences in the expression levels of *clpX* and *clpP* between the WT and the Δ*tssB* background at any of the tested time points.

The ClpXP degradation system recognizes its substrates via the C-terminal region, for instance for proteins that were tagged by the SsrA system (Sauer & Baker, 2011), and binds it within the axial pore of the ClpX ATPase (Martin *et al*, 2008), facilitating the enzyme’s ability to unfold substrates and translocate polypeptides into ClpP for degradation (Wawrzynow *et al*, 1995; Wojtkowiak *et al*, 1993). The canonical sequence of the SsrA-tag, consisting of 11 residues (AADENYNYALAA), is recognized by ClpX at the last three C-terminal amino acids (Flynn *et al*, 2001). Interestingly, when represented as a hexamer, the crystal structure of the Hcp protein from *A. baumannii* strain AB0057 (Ruiz *et al*, 2015) shows that its C-terminal domain is exposed, potentially making it accessible for interaction with ClpX and subsequent degradation (Fig. S3F). Additionally, the last 11 C-terminal residues of the *A. baumannii* A118 Hcp protein are SLSNNTASYAA. Thus, one can speculate that when Hcp fails to be secreted and accumulates, this ‘SsrA-like’ tag might be recognized by the ClpXP protease machinery, leading to degradation. However, under secretion-permissive conditions, the Hcp hexamer is enclosed within the contractile sheath, thereby hiding the SsrA-like tag and protecting Hcp from degradation. These observations suggest that *A. baumannii* has evolved a sophisticated regulation of its T6SS, closely linked to the strain’s secretion capacity to prevent unnecessary protein accumulation.

## Conclusion

In conclusion, our study reveals a novel role of the capsular polysaccharide in *A. baumannii*, highlighting its complex interaction with the T6SS, which is crucial for environmental colonization and survival. Notably, we demonstrated that both the capsule and the T6SS independently offer protection during antagonistic interactions with competitors, and these protective effects might be synergistic under laboratory conditions. However, overproduction of the capsule in *A. baumannii* A118 impedes T6SS activity by hindering its assembly, potentially due to increased membrane tension from excess polysaccharides affecting membrane complex anchoring. This inhibition is alleviated in the double mutant Δ*bfmS*Δ*itrA*, which does not produce the capsule, indicating that the polysaccharide directly inhibits T6SS. Furthermore, capsule overproduction disrupts the organization of polysaccharides on the cell surface, likely impairing proper T6SS secretion. While the inhibition observed in Δ*bfmS* could result from both increased membrane tension and altered surface organization, our data conclusively show that strains lacking capsules exhibit enhanced T6SS activity and secretion compared to their encapsulated counterparts, suggesting that the capsule imposes steric hindrance on T6SS.

Interestingly, the inhibition of T6SS activity that we observed in the capsule-overexpressing mutant (Δ*bfmS*) also occurs under conditions of antibiotic-induced capsule overexpression. It is plausible that this inhibition is an adaptive response to withstand antibiotic exposure. Ultimately, this trade-off between T6SS functionality and capsule-mediated protection poses a competitive disadvantage, with the optimal balance achieved in the wild type, where both systems are fully functional.

Furthermore, we found that in secretion-impaired strains, the accumulation of Hcp is mitigated by a degradation mechanism involving the general ClpXP protease machinery. Given that the production of the T6SS involves the continuous synthesis and secretion of hundreds of protein components, this degradation could serve as a strategy to alleviate proteotoxic stress and conserve energy, particularly under unfavorable conditions such as antibiotic presence.

In summary, this work establishes a foundational understanding of the interplay between extracellular polysaccharides, such as the capsule, and secretion processes in *A. baumannii*. This interaction urges further characterization to develop effective strategies against this problematic and often antibiotic-resistant pathogen.

## Materials and Methods

### Bacterial strains, plasmids, and growth conditions

The bacterial strains and plasmids utilized in this study are detailed in Appendix Table S1. Generally, bacteria were grown in lysogeny broth (LB-Miller; Carl Roth, Switzerland) or on LB agar plates, aerobically at 37°C. *E. coli* strains S17-1λpir (Simon *et al*, 1983) and MFDpir (Ferrières *et al*, 2010) served for cloning or as donors in mating experiments. For induction of the *P_BAD_* promoter, L-arabinose was added to the medium at a final concentration of 2%. Antibiotics and supplements were added as needed: kanamycin (50 μg/ml), carbenicillin (100 μg/ml), streptomycin (100 μg/ml), apramycin (100 μg/ml), chloramphenicol (5 μg/ml), gentamicin (15 μg/ml), diaminopimelic acid (0.3 mM DAP), and isopropyl β-D-1-thiogalactopyranoside (2 mM IPTG).

### Genetic engineering of *A. baumannii*

DNA manipulations adhered to established molecular biology protocols, using enzymes as per manufacturers’ directions. Enzymes were purchased from these companies: High-fidelity Q5 polymerase (New England Biolabs), GoTaq polymerase (Promega), T4 DNA ligase (New England Biolabs), and restriction enzymes (New England Biolabs). Engineered strains and plasmids underwent initial PCR screening and were finally validated by Sanger sequencing of PCR-amplified fragments or plasmids.

*A. baumannii* mutants were created via allelic exchange with the counter-selectable suicide plasmid pGP704-Sac-Kan (Metzger *et al*, 2019; Vesel & Blokesch, 2021). Briefly, deletion constructs or *msfGFP* gene fusions were crafted to include > 800 bp up– and downstream the target gene. These segments were amplified via PCR using oligonucleotides with 5’ restriction sites for later digestion. After digestion, the fragments were ligated with the similarly cut pGP704-Sac-Kan plasmid using T4 DNA ligase and then introduced into chemically competent *E. coli* S17-1λpir cells for further processes. Transformants were confirmed via colony PCR, and plasmid accuracy was ensured through Sanger sequencing. These plasmids were then introduced into *A. baumannii* through biparental mating for 8 h at 37°C. Selection of single-crossover transconjugants utilized CHROMagar *Acinetobacter* (CHROMagar, France) plates or LB agar with chloramphenicol and kanamycin. After mating, the transconjugants were incubated at 37°C for 16 h and then underwent selection at room temperature for the SacB-containing suicide plasmid’s loss using plates of NaCl-free LB agar containing 10% sucrose. Colony checks for antibiotic sensitivity confirmed plasmid loss. Mutants were validated through colony PCR and Sanger sequencing.

Selective mutants with antibiotic resistance markers were created via natural transformation, a method detailed in prior studies (Godeux *et al*, 2020; Vesel *et al*, 2023). The transforming material, generated by overlapping PCR, included a kanamycin resistance cassette (*aph*) flanked by FRT sites and 800 bp of homologous regions, enabling efficient transformation. Selected transformants on LB agar with kanamycin underwent verification through colony PCR and Sanger sequencing. The resistance cassette was then excised using the FLP/FRT recombinase system (Tucker *et al*, 2014), with the process and loss of the recombinase plasmid confirmed by antibiotic resistance tests, colony PCR, and Sanger sequencing, ensuring precise genetic manipulations.

### Interbacterial killing

The interbacterial killing assay was slightly modified from prior work (Flaugnatti *et al*., 2021). Bacteria were incubated overnight at 37°C with continuous shaking. They were then diluted 1:100 in fresh LB medium and grown until the optical density at 600nm (OD_600_) reached 1. For stationary-phase samples, overnight cultures after 15-16 h of growth were used directly. Bacterial cultures (1 ml) were concentrated to an OD_600_ of 10 with sterile PBS buffer. Predators and prey were mixed in 1:1 or 1:5 ratios and spotted onto filters placed on LB agar plates. After incubation at 37°C for 4 h, bacteria were resuspended in PBS, serially diluted, and spotted on selective media for an overnight incubation at 37°C. *A. baumannii* was selected on CHROMagar *Acinetobacter* medium (CHROMagar, France), while *E. coli* cells were selected on LB agar supplemented with streptomycin. Recovered colonies were counted to calculate the number of colony-forming units (CFU) per ml.

In this interbacterial killing assay to stimulate capsule production via chloramphenicol, bacteria were initially grown in LB for 20 h at 37°C, then 1:100 diluted and grown further for 20 h in LB without or with chloramphenicol (20 μg/ml). The cultures were subsequently processed as outlined above and the mixture of predators and treated prey was spotted onto LB agar, with or without chloramphenicol (25 μg/ml), and incubated at 37°C for 4h. After incubation, bacteria were resuspended in PBS, diluted, and plated on selective media for overnight growth at 37°C, as mentioned above. Each experiment was repeated three independent times. Statistical significance was determined based on log-transformed data, with detection limits defined by the absence of at least one recoverable prey bacterium.

### Hcp secretion assay

To assess Hcp secretion, bacteria were grown in LB medium overnight at 37°C, followed by a 1:100 dilution and further aerobic cultivation until reaching an OD_600_ of 1. For stationary-phase studies, overnight growth was extended to 15-16 h before proceeding with further analyses.

Chloramphenicol-treated samples underwent a similar initial growth phase for 20 h, followed by additional growth in the presence of chloramphenicol (20 μg/ml) for 20 h, maintaining the same aerobic growth conditions. Subsequently, 2 ml of the culture was collected through centrifugation (5 min, 8000 rpm) and the supernatant filtered (0.22-μm filter; VWR). Secreted proteins in the supernatant were then precipitated using 10% ice cold trichloroacetic acid (TCA) on ice for 2 h. To verify consistent precipitation, BSA (100 μg/ml) was added to the supernatant before precipitation. The precipitated proteins were washed with 100% acetone, resuspended in 2X Laemmli buffer (50 μl/OD unit of initial culture), and heated before undergoing SDS-PAGE and Western blot analysis.

### SDS-PAGE and Western blotting

Proteins were separated on 12% mini-protean TGX stain-free precast gels and transferred to a PVDF membrane using the Trans-blot system as per manufacturer’s instructions (Bio-Rad). Membranes were blocked in 2.5% skim milk at room temperature for 30 min. Primary antibodies were raised in rabbits against synthetic peptides of Hcp (Eurogentec) and used at a dilution of 1:667 in 2.5% skim milk. After 1.5 h of incubation, the membranes were washed three times with TBST (Tris-Buffered Saline with 0.1% Tween-20) buffer. They were then incubated for 1h with an anti-rabbit IgG conjugated to horseradish peroxidase (HRP) (A9169; Sigma-Aldrich) as the secondary antibody at a dilution of 1:10,000. Following three additional washes, the membranes were treated with Lumi-Light^PLUS^ Western Blotting substrate (Roche, Switzerland) for signal development and visualized using a ChemiDoc XRS+ station (Bio-Rad). The anti-Sigma70-HRP antibodies (BioLegend, USA distributed via Brunschwig, Switzerland) were used at a dilution of 1:10,000 to serve as a loading control in the experiment. Precipitated BSA was identified with anti-BSA-HRP-conjugated antibodies (Santa Cruz Biotechnology Inc.), diluted at 1:2,000, to verify the precipitation efficiency.

TssB-GFP fusions were identified using anti-GFP mouse monoclonal antibodies (1181446001; Roche) at a 1:5,000 dilution, with HRP-conjugated anti-mouse antibodies (A6782; Sigma-Aldrich) at 1:20,000 as secondary antibodies for 1 h.

### Serum killing assay

Bacteria were cultured overnight in 3 mL of LB medium under aerobic conditions at 37°C, then diluted 1:100 in fresh LB (2 ml) until the OD_600_ reached 1. Following centrifugation, 1 mL of culture was washed and resuspended in PBS adjusted to an OD_600_ of 1. For the assay, 40 μl of this bacterial suspension was mixed with 60 μL of baby rabbit complement serum (AbD Serotec) and incubated for 1 h at 37°C. Controls included PBS and heat-inactivated serum (heated at 56°C 30 minutes) treatments. The reactions were stopped by cooling the samples on ice, and surviving bacteria were quantified by plating serial dilutions on LB agar plates, incubated overnight at 37°C.

### Epifluorescence microscopy

After growing bacteria aerobically in 2 ml LB medium at 37°C to an OD_600_ of 1, they were applied to agarose pads (1% agarose dissolved in 1x PBS) mounted on glass slides and covered by a coverslip. Cell visualization was performed using a Zeiss LSM 700 inverted confocal microscope (Zeiss, Switzerland) equipped with a fluorescence light source (Illuminator HXP 120), an AxioCam MRm high resolution camera, and controlled by the Zeiss Zen software (ZEN blue edition). Image analysis was conducted with Fiji software (2.0.0-rc-69/1.53f/Java 1.8.0_202 (64-bit); (Schindelin *et al*, 2012)). The displayed images are representative of three independent biological replicates.

### Quantification of T6SS sheath structures

In the pre-processing stage, acquired images were corrected for both drift and photobleaching. Drift was adjusted using the linear stack alignment with the SIFT plugin in ImageJ, based on phase contrast images (Lowe, 2004). To compensate for photobleaching, a histogram matching method was applied to the fluorescence channel (Miura, 2020).

For sheath structure detection, the pre-processed fluorescence images were analyzed using ilastik software (Berg *et al*, 2019) to create two types of classifiers: a pixel classifier for identifying T6SS-positive pixels and an object classifier for categorizing T6SS objects as either dotted shaped (contracted) or rod-shaped (extended). These classifiers were made by manually annotating a set of images representative of the dataset variability. However, due to challenges in differentiating between contracted and extended sheath structures, this distinction was not made in the final analysis.

For bacterial segmentation, since they remain stationary and unchanged in shape throughout the acquisition, the first phase contrast time-point was utilized. The segmentation method has been previously published (Proutière *et al*, 2023).

Data analysis involved pre-processing, segmentation, and sheath structure classification steps performed in ImageJ/Fiji using a Groovy script for batch processing (WorkFlow_File.groovy). This script generated a new multi-channel time-lapse stack per image, which consisted of the drift-compensated phase contrast channel, the bleach and drift compensated fluorescent channel, a color-coded mask channel for contracted and extended sheath structures, and a label image of detected bacteria. The resulting stacks were used for visual assessment of the method and for downstream data analysis. A second script (CountObject_File) quantified the sheath structures per bacterium, per time point, and for each condition, output the data in a tsv file. All scripts, models, and classifiers were deposited on Zenodo (see data availability section below).

### Extraction of capsular polysaccharides

Polysaccharide samples from cell lysates (membrane-bound) and culture supernatants (membrane-unbound) were prepared using a procedure slightly modified from previous studies (Geisinger & Isberg, 2015; Tipton & Rather, 2019). Briefly, bacteria grown overnight on LB agar plates at 37°C were resuspended in LB medium and adjusted to an OD_600_ of 10. Cells were separated from the supernatant by centrifugation. The supernatant was then precipitated with 75% ethanol at –20°C overnight. Meanwhile, the cell fraction was resuspended in lysis buffer (60 mM Tris, pH 8, 10 mM MgCl_2_, 50 µM CaCl_2_ with 20 µl/ml DNase and 3 mg/mL lysozyme) and incubated at 37°C for 1 h. Post-vortexing, the cell fraction underwent three freeze-thaw cycles between liquid nitrogen and 37°C. The suspension was treated with 0.5% SDS for 30 min at 37°C, boiled at 100°C for 10 min, then treated with proteinase K (2 mg/ml) at 60°C for 1 h. Following centrifugation (2 min, 15000 x *g*), the supernatant was precipitated with 75% ethanol at –20°C overnight. The precipitated polysaccharides were centrifuged (30 min, 15000 x *g*), resuspended in 40 μl 2X Laemmli buffer (Sigma-Aldrich, Switzerland), heated at 95°C for 10 min, and analyzed by SDS-PAGE and Alcian blue staining.

Samples treated with chloramphenicol were grown for 20 h in LB medium, followed by additional growth in LB medium supplemented with varying concentrations of chloramphenicol (0, 2.5, 5, 7.5, 10, or 20 µg/ml). Polysaccharides from the culture supernatants were precipitated and processed as explained above.

### Alcian blue staining

Precipitated polysaccharide samples were run on 12% mini-protean TGX stain-free precast gels (Bio-Rad) and stained with Alcian blue (0.1% Alcian blue in 40% ethanol, 60% 20 mM sodium acetate (pH 4.75)) for 1 h, then destained overnight (40% ethanol, 60% 20 mM sodium acetate (pH 4.75)) (Karlyshev & Wren, 2001).

### Capsule visualization by Transmission Electron Microscopy (TEM)

For TEM visualization of capsular polysaccharide, previously published protocols were followed with minor modifications (Chin *et al*., 2018; Valcek *et al*, 2023). Briefly, the bacterial strains were cultured in LB medium for 15-16 h at 37°C under shaking conditions, then a 500 μl sample was centrifuged to form a small pellet. This pellet was fixed on ice for 20 min with a mixture containing 2% paraformaldehyde and 2.5% glutaraldehyde in 0.1M sodium cacodylate buffer (pH 7.4) with 1.55% L-lysine acetate and 0.075 % ruthenium red. Following this, the fixed bacteria were washed three times in 0.1M sodium cacodylate buffer (pH 7.4) with 0.075% ruthenium red. A second round of fixation was performed in the same fixation solution minus the lysine acetate for 2h. This fixation was followed by two additional rounds of washing in sodium cacodylate/ruthenium red buffer and then staining with 1% osmium tetroxide and 0.075% ruthenium red in 0.1M cacodylate buffer for 1h at room temperature. Finally, the sample was washed with 0.075% ruthenium red in 0.1 M cacodylate buffer followed by distilled water and then dehydrated in a graded ethanol series before being embedded in an epon resin (Embed 812 embedding kit, EMS), which was polymerized for 24h at 60°C.

After hardening, 50 nm sections were prepared with a Leica UC7 Ultramicrotome and collected onto single-slot copper grids with a pioloform support film. The sections underwent contrasting for enhanced visibility with 2% lead citrate and 1% uranyl acetate and were imaged using a TEM (FEI Spirit) with a CCD camera (FEI Eagle) to capture each cell’s structure.

### Structural model of the Hcp hexamer

The structural model of the Hcp hexamer is based on the crystal structure of the Hcp1 protein of *A. baumannii* strain AB0057 (RCSB PDB (Berman *et al*, 2000) code 4W64 (Ruiz *et al*., 2015)) and was visualized using Mol* Viewer 2.9.3 (Sehnal *et al*, 2021).

## Data availability

- Imaging dataset: All scripts, models, and classifiers relevant to the image analyses have been deposited on Zenodo (doi: 10.5281/zenodo.11039744).
- All other data are part of the manuscript or the supplementary material.

## Acknowledgments

The authors would be like to acknowledge current and former members of the Blokesch group for insightful discussions, with special thanks to Nina Vesel for advice on *A. baumannii* genetics and David W. Adams for general scientific advice, and Charles Van der Henst, Graham Knott, and Christel Genoud for helpful suggestions on Transmission Electron Microscopy. The authors also acknowledge the technical assistance of Sandrine Stutzmann, Laurie Righi, and Candice Stoudmann, and recognize Lisa Metzger for plasmid engineering. The authors thank Nicolas Chiaruttini from the BioImaging and Optics platform of EPFL for development of the T6SS quantification script. The authors are grateful to Marek Basler, Xavier Charpentier, and Bryan W. Davies for sharing of *A. baumannii* strains or plasmids. This work was supported by the Swiss National Science Foundation (grant numbers 407240_167061 and 310030_204335) and a Howard Hughes Medical Institute (HHMI) International Research Scholarship (grant number 55008726) attributed to M.B.

## Author contributions

M.B. supervised the work and secured funding; M.B and N.F. conceived the project and analyzed the results; N.F. planned the experiments; N.F., L.B., M.C.-C. performed the experiments; N.F. and M.B. wrote the manuscript. All authors approved the final version of the manuscript.

## Author’s ORCID numbers

Nicolas Flaugnatti https://orcid.org/0000-0002-6073-3340

Melanie Blokesch https://orcid.org/0000-0002-7024-1489

## Supplementary Figure legends

**Figure S1: *A. baumannii* strain A118 produces functional T6SS**

**(A)** Illustration of the T6SS gene arrangement in *A. baumannii* strain A118. The core components of the T6SS are highlighted in gray. **(B)** Survival of *E. coli* following interaction with WT (strain A118), the two T6SS-inactive mutants (Δ*hcp* and Δ*tssB*), or the TssB translational fusion-carrying strain (*tssB*-*msfGFP*). Survival rates are presented on the *Y*-axis. Data points are from three independent experiments, with bars indicating mean values (± SD, depicted by error bars). Statistical significance was determined using an ordinary one-way ANOVA test. \*\*\*\**P* < 0.0001, ns = not significant. The detection limit (dl) is indicated.

**Figure S2: Deletion of *bfmS* and its effect on T6SS activity in *A. baumannii***

**(A)** and (**B**) Colony morphologies on blood agar plates after 24h of growth, with strain genotypes indicated. (**B**) depicts zoomed regions of the white boxes shown in panel A. **(C)** Complementation of *bfmS* deletion assessed by T6SS activity. Enumeration of *E. coli* after exposure to WT, T6SS-inactive (Δ*hcp*), Δ*bfmS*, and the *bfmS*-complemented strain (Δ*bfmS*::*bfmS*), with survival shown on the *Y*-axis. **(D)** Hcp secretion remains detectable in WT co-cultured with secretion-impaired (+Δ*tssB*) or capsule-overproducing Δ*bfmS* (+Δ*bfmS*) strains, as analyzed by immunoblotting. Details as described in Figure 2B. **(E)** Survival of *E. coli* prey after contact with *A. baumannii* WT and mutants Δ*hcp,* Δ*bfmS* strains across various strain backgrounds (A118, 29D2, and 86II/2C). Details as in panel C. **(F)** Equal TssB production in varies strains. TssB production was assessed in exponentially growing strains carrying a translational fusion of the T6SS sheath protein TssB and msfGFP (*tssB-msfGFP*) by immunoblot analysis using anti-GFP antibodies. Strain backgrounds: WT, Δ*itrA*, Δ*bfmS*, and Δ*bfmS*Δ*itrA*. Equal loading of the cell lysates (CL) was confirmed by detection of σ70.

**Figure S3: Hcp degradation is conserved across *A. baumannii* strains**.

**(A)** Hcp levels in WT and the Δ*tssB* mutant of *A. baumannii* strains A118, 29D2 and 86IIC were evaluated under exponential (left) and stationary (right) growth phases via immunoblot analysis. Details as described for Figure 2B. **(B)** TssB production remains equal during stationary phase. Strains harboring the translational fusion TssB-msfGFP were cultured under stationary growth conditions and analyzed for GFP production. Details on strains and immunoblotting conditions as described for panel S2F. **(C)** Growth curve of WT and the Δ*tssB* mutant over a 16-hour timeframe, with the gray zone highlighting the observed period of Hcp degradation shown in Figure 6C. (**D**) Assessment of *hcp*-overexpression plasmid in *E. coli*. The graph shows relative *hcp* expression levels in *E. coli* with an empty plasmid (+p) versus those with a plasmid for *hcp* overexpression (+p-*hcp*). The images below the graph illustrate Hcp production in these *E. coli* strains, which were assessed by immunoblotting. **(E)** Relative expression of *clpX* and *clpP* over time. The panel compares the relative expression of *clpX* and *clpP* over a 12-hour period in WT and Δ*tssB* strains. **(F)** The C-terminus of Hcp is surface-exposed. The presented Hcp hexamer is based on PDB 4W64 (Ruiz *et al*., 2015) with the C-termini color-coded in pink. Data are representative of three independent experiments. For the graphs in panels (**D**) and (**E**), data are represented as means (± SD, as indicated by error bars). Statistical significance was assessed using a two-way ANOVA on log-transformed data, comparing values between the vector control (+p) or plasmid p-*hcp* (**B**) or between the WT and Δ*tssB* conditions at 2 h versus later timepoints (**E**). \*\**P* < 0.01. Statistical values showing no significant differences have been omitted for clarity.

**Movie S1: Movie depicting image analysis pipeline**.

A representative movie is shown; snapshots were taken every 30 seconds over 5 minutes. The panel displays split views: the phase contrast channel on the left, the bleach and drift-compensated fluorescent channel in the middle, and the color-coded channel for contracted and extended sheath structures on the right. A mask depicting the segmented bacteria is overlaid in all three panels.

