## Supplemental Material for "Capsular Polysaccharide Restrains Type VI Secretion in *Acinetobacter baumannii*"

**<u>Table of content</u>**

**Figure S1:** ***A. baumannii*** **strain A118 produces functional T6SS**

**Figure S2: Deletion of** ***bfmS*** **and its effect on T6SS activity in** ***A. baumannii***

**Figure S3: Hcp degradation is conserved across** ***A. baumannii*** **strains.**

**Movie S1: Movie depicting image analysis pipeline.** [separate file]

**Appendix Table S1. Strains and plasmids used on this study.** [separate file]

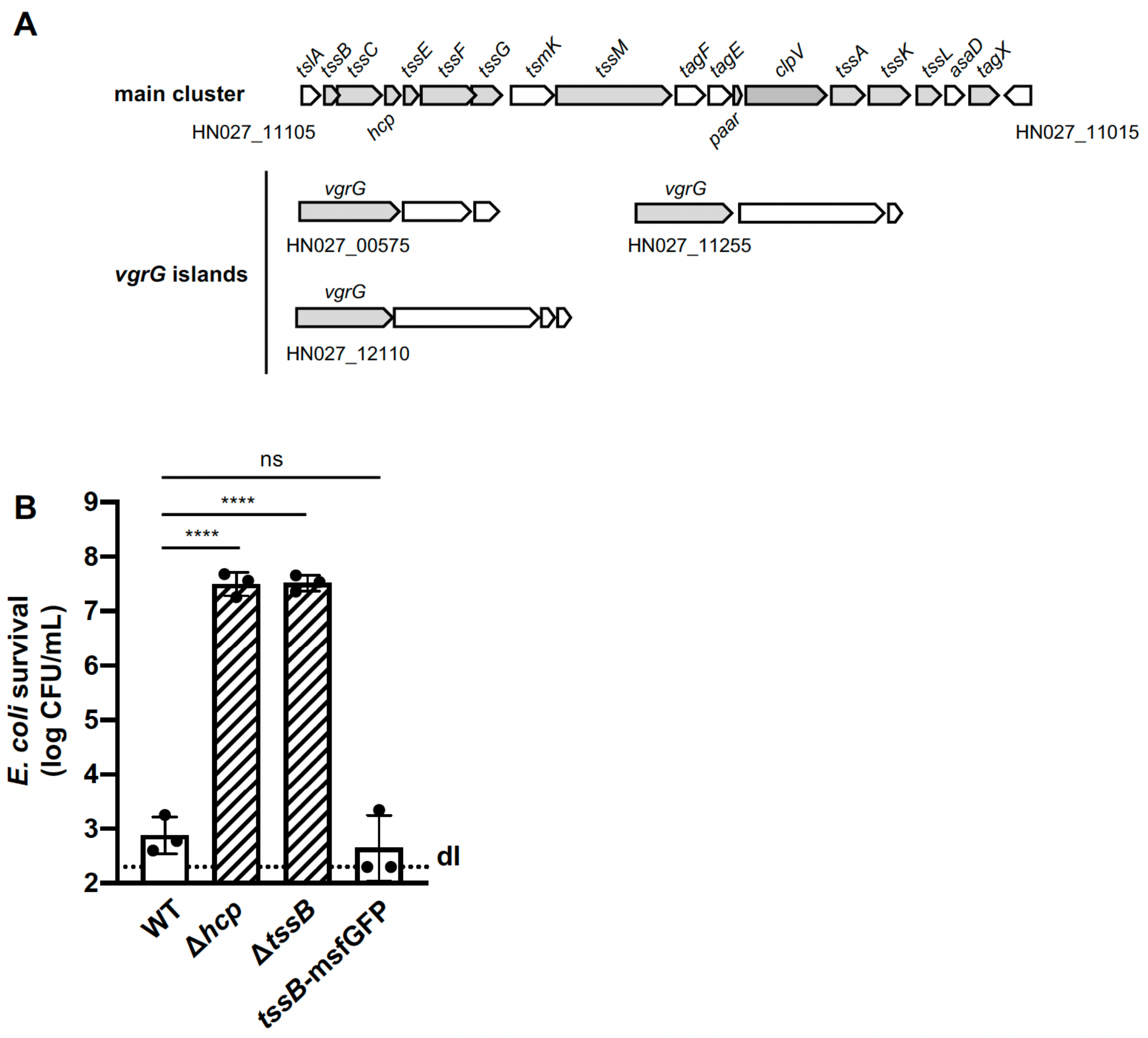


**Figure S1:** ***A. baumannii*** **strain A118 produces functional T6SS**
**(A)** Illustration of the T6SS gene arrangement in *A. baumannii* strain A118. The core components of the T6SS are highlighted in gray. **(B)** Survival of *E. coli* following interaction with WT (strain A118), the two T6SS-inactive mutants (Δ*hcp* and Δ*tssB*), or the TssB translational fusion-carrying strain (*tssB-msfGFP*). Survival rates are presented on the *Y*-axis. Data points are from three independent experiments, with bars indicating mean values (± SD, depicted by error bars). Statistical significance was determined using an ordinary one-way ANOVA test. ****$P < 0.0001$, ns = not significant. The detection limit (dl) is indicated.

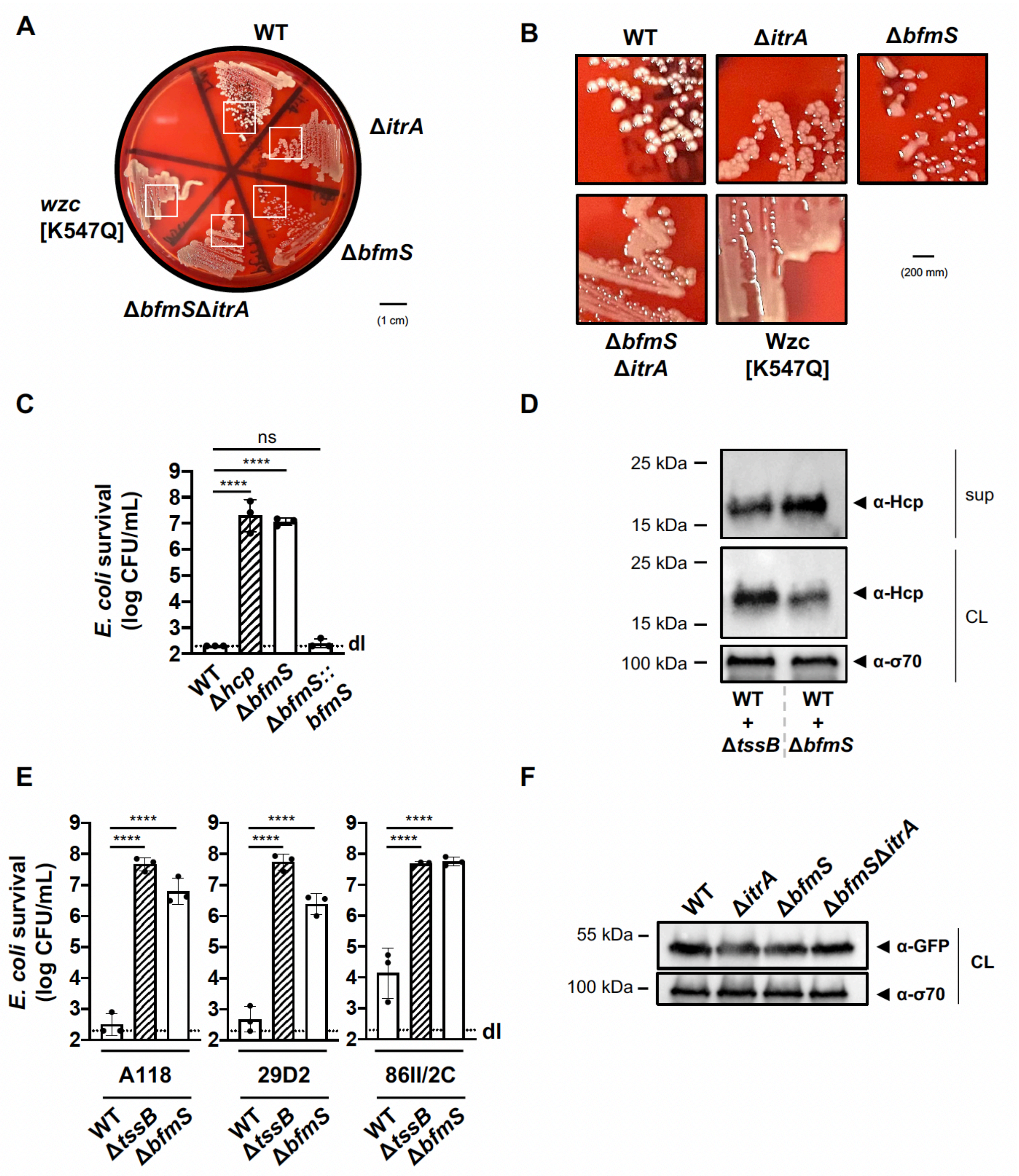


**Figure S2: Deletion of *bfmS* and its effect on T6SS activity in *A. baumannii***

**(A)** and **(B)** Colony morphologies on blood agar plates after 24h of growth, with strain genotypes indicated. **(B)** depicts zoomed regions of the white boxes shown in panel A. **(C)** Complementation of *bfmS* deletion assessed by T6SS activity. Enumeration of *E. coli* after exposure to WT, T6SS-inactive (Δ*hcp*), Δ*bfmS*, and the *bfmS*-complemented strain (Δ*bfmS*::*bfmS*), with survival shown on the *Y*-axis. **(D)** Hcp secretion remains detectable in WT co-cultured with secretion-impaired (+Δ*tssB*) or capsule-overproducing Δ*bfmS* (+Δ*bfmS*) strains, as analyzed by immunoblotting. Details as described in Figure 2B. **(E)** Survival of *E. coli* prey after contact with *A. baumannii* WT and mutants Δ*hcp,* Δ*bfmS* strains across various strain backgrounds (A118, 29D2, and 86II/2C). Details as in panel C. **(F)** Equal TssB production in varies strains. TssB production was assessed in exponentially growing strains carrying a translational fusion of the T6SS sheath protein TssB and msfGFP (*tssB-msfGFP*) by immunoblot analysis using anti-GFP antibodies. Strain backgrounds: WT, Δ*itrA* , Δ*bfmS*, and Δ*bfmS*Δ*itrA*. Equal loading of the cell lysates (CL) was confirmed by detection of σ70.

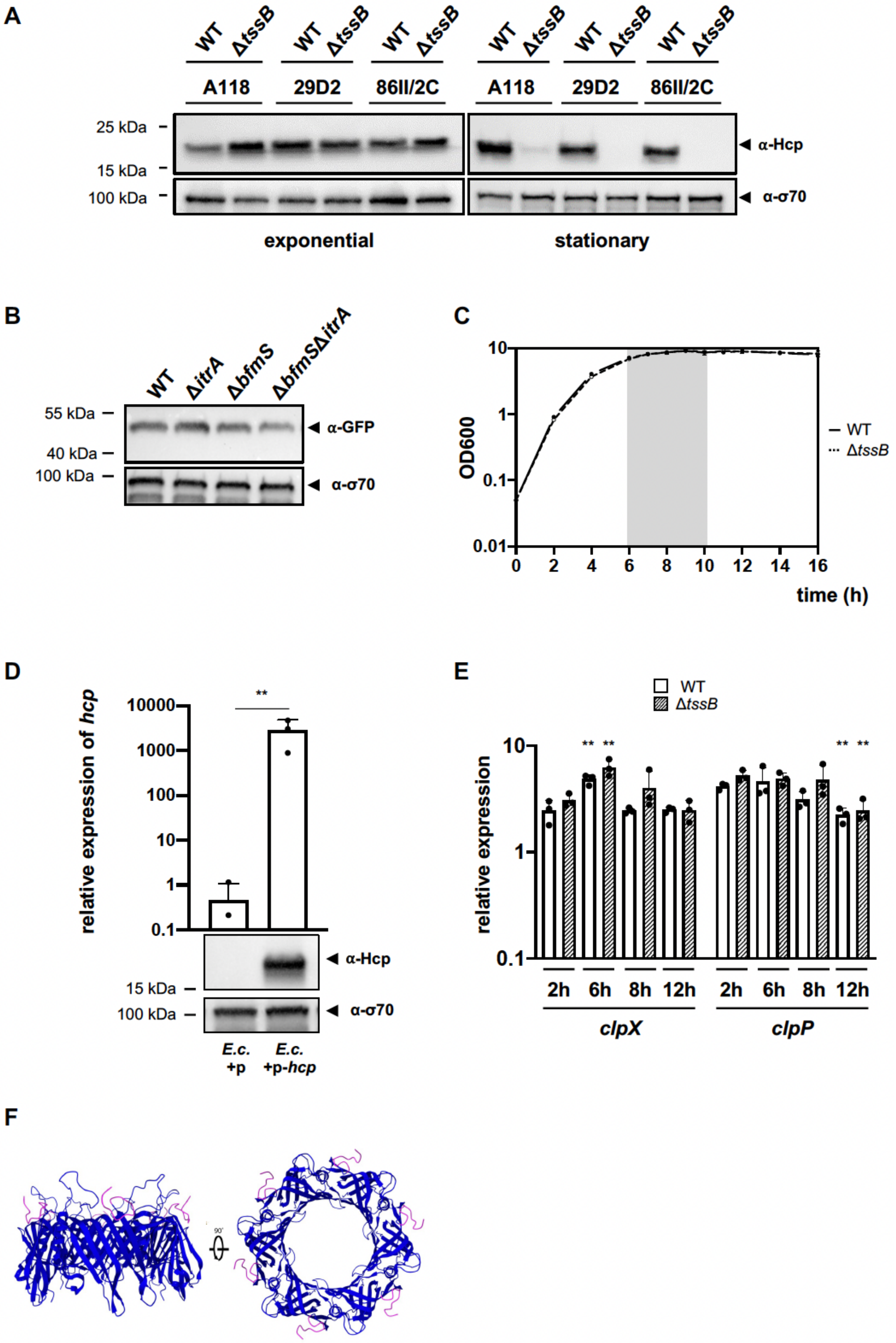

A
WT ΔtssB WT ΔtssB WT ΔtssB WT ΔtssB WT ΔtssB WT ΔtssB
A118 29D2 86II/2C A118 29D2 86II/2C
25 kDa
15 kDa
100 kDa
α-Hcp
α-σ70
exponential
stationary
B
WT ΔitrA ΔbfmS ΔbfmSΔitrA
55 kDa
40 kDa
100 kDa
α-GFP
α-σ70
C
OD600
10
1
0.1
0.01
0 2 4 6 8 10 12 14 16
time (h)
WT
ΔtssB
D
relative expression of hcp
10000
1000
100
10
1
0.1
**
α-Hcp
α-σ70
15 kDa
100 kDa
E.c. +p
E.c. +p-hcp
E
WT
ΔtssB
relative expression
10
1
0.1
** ** ** **
2h 6h 8h 12h 2h 6h 8h 12h
clpX
clpP
F
90°

**Figure S3: Hcp degradation is conserved across *A. baumannii* strains.**

**(A)** Hcp levels in WT and the Δ*tssB* mutant of *A. baumannii* strains A118, 29D2 and 86IIC were evaluated under exponential (left) and stationary (right) growth phases via immunoblot analysis. Details as described for Figure 2B. **(B)** TssB production remains equal during stationary phase. Strains harboring the translational fusion TssB-msfGFP were cultured under stationary growth conditions and analyzed for GFP production. Details on strains and immunoblotting conditions as described for panel S2F. **(C)** Growth curve of WT and the Δ*tssB* mutant over a 16-hour timeframe, with the gray zone highlighting the observed period of Hcp degradation shown in Figure 6C. (**D**) Assessment of *hcp*-overexpression plasmid in *E. coli*. The graph shows relative *hcp* expression levels in *E. coli* with an empty plasmid (+p) versus those with a plasmid for *hcp* overexpression (+p-*hcp*). The images below the graph illustrate Hcp production in these *E. coli* strains, which were assessed by immunoblotting. **(E)** Relative expression of *clpX* and *clpP* over time. The panel compares the relative expression of *clpX* and *clpP* over a 12-hour period in WT and Δ*tssB* strains. **(F)** The C-terminus of Hcp is surface-exposed. The presented Hcp hexamer is based on PDB 4W64 (Ruiz *et al.*, 2015) with the C-termini color-coded in pink. Data are representative of three independent experiments. For the graphs in panels (**D**) and (**E**), data are represented as means (± SD, as indicated by error bars). Statistical significance was assessed using a two-way ANOVA on log-transformed data, comparing values between the vector control (+p) or plasmid p-*hcp* (**B**) or between the WT and Δ*tssB* conditions at 2 h versus later timepoints (**E**). $^{**}P < 0.01$. Statistical values showing no significant differences have been omitted for clarity.
