## Supplementary material for "Capsular Polysaccharide Restrains Type VI Secretion in *Acinetobacter baumannii*": Table S1

### Appendix Table S1. Strains and plasmids used on this study.

| Strains or plasmids | Genotype / description | Strain number | Source and/or reference |
|---|---|---|---|
| ***Acinetobacter baumannii*** | | | |
| A118 | Wild type; $Amp^R$, $Cm^R$; ATCC BAA-2093 | MB#5144 | ATCC (ATCC BAA-2093) (Ramirez *et al*, 2010) (Traglia *et al*, 2014) |
| A118Δ*itrA* | A118 with *itrA* deleted using suicide plasmid pGP704-Sac-Kan-Δ*itrA* | MB#11161 | This study |
| A118Δ*hcp* | A118 with *hcp* deleted using suicide plasmid pGP704-Sac-Kan-Δ*hcp* | MB#11160 | This study |
| A118Δ*hcp*Δ*itrA* | A118Δ*hcp* with *itrA* deleted using suicide plasmid pGP704-Sac-Kan-Δ*itrA* | MB#11162 | This study |
| A118Δ*hcp*-TnAraC | A118Δ*hcp* containing mini-Tn7-*araC* (TnAraC); $Cm^R$, $Gent^R$ | MB#11165 | This study |
| A118Δ*hcp*Δ*itrA*-TnAraC | A118Δ*hcp*Δ*itrA* containing mini-Tn7-*araC* (TnAraC); $Cm^R$, $Gent^R$ | MB#11164 | This study |
| A118Δ*hcp*Δ*itrA*-Tn*itrA* | A118Δ*hcp*Δ*itrA* containing mini-Tn7-*araC-itrA* (Tn*itrA*); $Cm^R$, $Gent^R$ | MB#11163 | This study |
| A118-*tssB-msfgfp* | A118 carrying translational fusion encoding *tssB-msfgfp* at native *tssB* locus; constructed using suicide plasmid pGP704-Sac-Kan-*tssB-msfgfp* | MB#11201 | This study (fusion derived from Lin *et al.*, 2022) |
| A118Δ*itrA-tssB-msfgfp* | A118 carrying translational fusion encoding *tssB-msfgfp* at native *tssB* locus with *itrA* deleted using suicide plasmid pGP704-Sac-Kan-Δ*itrA* | MB#11200 | This study |
| A118Δ*tssB*::FRT-kan-FRT2 | A118 with *tssB* deleted by natural transformation with PCR fragment replacing *tssB* by FRT-*aph*-FRT2 cassette; $Kan^R$ | MB#11199 | This study |
| A118Δ*tssB*::FRT | A118Δ*tssB*::FRT-kan-FRT2 after flip and cure | MB#11198 | This study |
| A118Δ*bfmS* | A118 with *bfmS* deleted using suicide plasmid pGP704-Sac-Kan-Δ*bfmS* | MB#11197 | This study |
| A118Δ*bfmS*Δ*itrA* | A118Δ*bfmS* with *itrA* deleted using suicide plasmid pGP704-Sac-Kan-Δ*itrA* | MB#11196 | This study |
| A118-*wzc*[K547Q] | A118 carrying a substitution in the Wzc walker A box [K547Q] made using suicide plasmid pGP704-Sac-Kan-*wzc*[K547Q] | MB#11195 | This study |
| A118-*glmS*-$P_{[bfmS]}$ | A118 transformed by natural transformation with PCR fragment inserted between *glmS* and *murI,* containing upstream intergenic region (77 bp) of *bfmS* followed by *aac(3)IV* cassette; $Apr^R$ | MB#11194 | This study |

| | | | |
|---|---|---|---|
| A118Δ*hcp-glmS-P*[*bfmS*] | A118Δ*hcp* transformed by natural transformation with PCR fragment inserted between *glmS* and *murI*, containing upstream intergenic region (77 bp) of *bfmS* followed by *aac(3)IV* cassette; $Apr^R$ | MB#11193 | This study |
| A118Δ*bfmS-glmS-P*[*bfmS*] | A118Δ*bfmS* transformed by natural transformation with PCR fragment inserted between *glmS* and *murI*, containing upstream intergenic region (77 bp) of *bfmS* followed by *aac(3)IV* cassette; $Apr^R$ | MB#11192 | This study |
| A118Δ*bfmS-glmS-P*[*bfmS*]*-bfmS* | A118Δ*bfmS* transformed by natural transformation with PCR fragment inserted between *glmS* and *murI*, containing upstream intergenic region (77 bp) of *bfmS* and *bfmS* followed by *aac(3)IV* cassette; $Apr^R$ | MB#11191 | This study |
| A118Δ*tslA-tssB-msfgfp* | A118 with *tslA* deleted carrying a translational fusion encoding *tssB-msfgfp* at native *tssB* locus using suicide plasmid pGP704-Sac-Kan-*tssB-msfgfp* | MB#11476 | This study |
| A118Δ*bfmS-tssB-msfgfp* | A118 with *bfmS* deleted carrying translational fusion encoding *tssB-msfgfp* at native *tssB* locus using suicide plasmid pGP704-Sac-Kan-*tssB-msfgfp* | MB#11188 | This study |
| A118Δ*bfmS*Δ*itrA-tssB-msfgfp* | A118Δ*bfmS* carrying a translational fusion encoding *tssB-msfgfp* at native *tssB* locus with *itrA* deleted using suicide plasmid pGP704-Sac-Kan-Δ*itrA* | MB#11187 | This study |
| A118Δ*tslA* | A118 with deleted *tslA* using suicide plasmid pGP704-Sac-Kan-Δ*tslA* (plasmid from (Lin *et al*, 2022), as listed below) | MB#11186 | This study |
| A118 / pMMB67EH | A118 carrying plasmid pMMB67EH; $Amp^R$ | MB#11174 | This study |
| A118Δ*hcp* / pMMB67EH | A118Δ*hcp* carrying plasmid pMMB67EH ; $Amp^R$ | MB#11172 | This study |
| A118 / pMMB67EH-*hcp* | A118 carrying plasmid pMMB67EH-*hcp*; $Amp^R$ | MB#11173 | This study |
| A118Δ*hcp* / pMMB67EH-*hcp* | A118Δ*hcp* carrying plasmid pMMB67EH-*hcp*; $Amp^R$ | MB#11171 | This study |
| A118Δ*clpXP*::FRT-kan-FRT2 | A118 with *clpX* and *clpP* deleted by natural transformation with PCR fragment containing *clpX* and *clpP* replaced by FRT-*aph*-FRT2 cassette; $Kan^R$ | MB#11170 | This study |
| A118Δ*lon*::FRT-kan-FRT2 | A118 deleted for *lon* by natural transformation with PCR fragment containing *lon* replaced by FRT-*aph*-FRT2 cassette; $Kan^R$ | MB#11179 | This study |
| A118Δ*tssB*::FRTΔ*clpXP*::FRT-kan-FRT2 | A118Δ*tssB*::FRT deleted for *clpX* and *clpP* genes by natural transformation with PCR fragment containing *clpX* and *clpP* replaced by FRT-*aph*-FRT2 cassette; $Kan^R$ | MB#11169 | This study |
| A118Δ*tssB*::FRTΔ*lon*::FRT-kan-FRT2 | A118Δ*tssB*::FRT deleted for *lon* by natural transformation with PCR fragment containing *lon* replaced by FRT-*aph*-FRT2 cassette; $Kan^R$ | MB#11178 | This study |
| 29D2 | Wild type; $Amp^R$ | MB#8581 | (Wilharm *et al*, 2017) |
| 29D2Δ*tssB::*FRT-kan-FRT2 | 29D2 deleted for *tssB* by natural transformation with PCR fragment containing *tssB* replaced by FRT-*aph*-FRT2 cassette; $Kan^R$ | MB#11177 | This study |
| 29D2Δ*bfmS* | 29D2 with *bfmS* deleted using suicide plasmid pGP704-Sac-Kan-Δ*bfmS* | MB#11168 | This study |
| 86II/2C | Wild type; $Amp^R$ | MB#8581 | (Wilharm *et al.*, 2017) |

| | | | |
|---|---|---|---|
| 86II/2Δ*tssB::*FRT-kan-FRT | 86II/2C deleted for *tssB* by natural transformation with PCR fragment containing *tssB* replaced by FRT-*aph*-FRT; $Kan^R$ | MB#11176 | This study |
| 86II/2CΔ*bfmS* | 86II/2C with *bfmS* deleted using suicide plasmid pGP704-Sac-Kan-Δ*bfmS* | MB#11167 | This study |
| ***Enterobacter cloacae*** | | | |
| ERR2221156 | *Enterobacter cloacae* commensal ERR2221156 from HBC collection | MB#8341 | (Forster *et al*, 2019) |
| ERR2221156Δ*tssB* | ERR2221156 deleted for *tssB* | MB#9173 | (Flaugnatti *et al*, 2021) |
| ***Escherichia coli* strains** | | | |
| S17-1λpir | $Tp^R$ $Sm^R$ *recA thi pro hsdR2M1* RP4:2-Tc:Mu:Kmr Tn7 (λ*pir*) | MB#648 | (Simon *et al*, 1983) |
| SM10λpir | *thi-1 thr leu ton*A *lac*Y *sup*E *rec*A::RP4-2-Tc::Mu, $Km^R$ (λ*pir*) | MB#647 | Laboratory stock |
| TOP10 | F- *mcrA* Δ(*mrr-hsd*RMS-*mcr*BC) φ80*lacZ*ΔM15 Δ*lac*X74 *nup*G *rec*A1 *ara*Δ139 Δ(*ara-leu*)7697 *gal*E15 *gal*K16 *rpsL* ($Strep^R$) *end*A1λ-. | MB#741 | Invitrogen |
| TOP10 / pMMB67EH | TOP10 containing pMMB67EH vector; $Amp^R$ | MB#11203 | This study |
| TOP10 / pMMB67EH-*hcp* | TOP10 containing plasmid pMMB67EH-*hcp*; $Amp^R$ | MB#11175 | This study |
| K-12 | F+ lambda+ K-12 strain | MB#2903 | Laboratory stock |
| K-12 / pSG3685 | *E. coli* K-12 transformed by electroporation with plasmid pSG3685 to confer $Strep^R$ | MB#11202 | This study |
| MC4100 | F- [*araD*139]B/r DE(*argF-lac*)169 Lambda- e14- *flh*D5301 DE(*fruK-yeiR*)725(*fruA25*) *relA1 rpsL150*($Strep^R$) rbsR22 DE(*fimB-fimE*)632(::IS1) deoC1 | MB#2905 | Laboratory stock |
| MC4100-Tn-CmR | *E. coli* MC4100 containing mini-Tn7-araC-*cat* (Tn-CmR) confering resistance to chloramphenicol; $Cm^R$, $Strep^R$ | MB#5123 | This study |
| **Plasmids** | | | |
| pGP704-Sac28 | Suicide plasmid; oriR6K *sacB*; $Amp^R$ | MB#649 | (Meibom *et al*, 2004) |
| pGP704-Sac-Kan | Suicide plasmid, oriR6K *sacB*; $Kan^R$ | MB#6038 | (Metzger *et al*, 2019) |
| pGP704-TnAraC | pGP704 with mini-Tn7 carrying *araC* and $P_{BAD}$; $Amp^R$, $Gent^R$ | MB#5513 | (Adams *et al*, 2019) |
| pUX-BF13 | oriR6K, helper plasmid with Tn7 transposition function; $Amp^R$ | MB#457 | (Bao *et al*, 1991) |
| pGP704-Sac-Kan-Δ*itrA* | pGP704-Sac-Kan carrying a deletion within *itrA*; $Kan^R$ | MB#11185 | This study |
| pGP704-Sac-Kan-Δ*hcp* | pGP704-Sac-Kan carrying a deletion within *hcp*; $Kan^R$ | MB#11184 | This study |

| | | | |
|---|---|---|---|
| pGP704-Sac-Kan-Δ*bfmS* | pGP704-Sac-Kan carrying a deletion within *bfmS*; $Kan^R$ | MB#11183 | This study |
| pGP704-Sac-Kan-*tssB-msfgfp* | pGP704-Sac-Kan carrying a translational fusion encoding *tssB-msfgfp* to replace *tssB* at its native locus; the insert was amplified using genomic DNA of strain LLB832 from Lin *et al.*, 2022 as template; $Kan^R$ | MB#11182 | This study (fusion derived from Lin *et al.*, 2022) |
| pGP704-Sac-Kan-*wzc*[K547Q] | pGP704-Sac-Kan encoding K547Q variant of Wzc; $Kan^R$ | MB#11181 | This study |
| pGP704-Sac-Kan-Δ*tslA* | pGP704-Sac-Kan carrying a deletion within *tslA*; $Kan^R$ | MB#11189 | (Lin *et al.*, 2022) |
| pGP704-Tn-ItrA | pGP704 with mini-Tn7 carrying *araC* and $P_{BAD}$-driven *itrA*; $Amp^R$, $Gent^R$ | MB#11180 | This study |
| pGP704-Tn-$Cm^R$ | pGP704 with mini-Tn7 carrying *cat* cassette; $Amp^R$, $Gent^R$, $Cm^R$ | MB#5054 | (Metzger *et al.*, 2019) |
| pAT03 | pMMB67EH with FLP recombinase gene; $Amp^R$ | MB#9372 | (Tucker *et al*, 2014) |
| pMMB67EH | pMMB67EH; $Amp^R$ | MB#9371 | Laboratory stock |
| pMMB67EH-*hcp* | pMMB67EH with *hcp* gene from *A. baumannii* A118; $Amp^R$ | MB#11175 | This study |

**References Appendix Table S1**


Adams DW, Stutzmann S, Stoudmann C, Blokesch M (2019) DNA-uptake pili of *Vibrio cholerae* are required for chitin colonization and capable of kin recognition via sequence-specific self-interaction. *Nat Microbiol* 4: 1545-1557

Bao Y, Lies DP, Fu H, Roberts GP (1991) An improved Tn*7*-based system for the single-copy insertion of cloned genes into chromosomes of Gram-negative bacteria. *Gene* 109: 167-168

Flaugnatti N, Isaac S, Lemos Rocha LF, Stutzmann S, Rendueles O, Stoudmann C, Vesel N, Garcia-Garcera M, Buffet A, Sana TG *et al* (2021) Human commensal gut Proteobacteria withstand type VI secretion attacks through immunity protein-independent mechanisms. *Nat Commun* 12: 5751

Forster SC, Kumar N, Anonye BO, Almeida A, Viciani E, Stares MD, Dunn M, Mkandawire TT, Zhu A, Shao Y *et al* (2019) A human gut bacterial genome and culture collection for improved metagenomic analyses. *Nat Biotechnol* 37: 186-192

Lin L, Capozzoli R, Ferrand A, Plum M, Vettiger A, Basler M (2022) Subcellular localization of Type VI secretion system assembly in response to cell-cell contact. *EMBO J* 41: e108595

Meibom KL, Li XB, Nielsen AT, Wu CY, Roseman S, Schoolnik GK (2004) The *Vibrio cholerae* chitin utilization program. *Proc Natl Acad Sci USA* 101: 2524-2529

Metzger LC, Matthey N, Stoudmann C, Collas EJ, Blokesch M (2019) Ecological implications of gene regulation by TfoX and TfoY among diverse *Vibrio* species. *Environ Microbiol* 21: 2231-2247

Ramirez MS, Don M, Merkier AK, Bistue AJ, Zorreguieta A, Centron D, Tolmasky ME (2010) Naturally competent *Acinetobacter baumannii* clinical isolate as a convenient model for genetic studies. *J Clin Microbiol* 48: 1488-1490

Simon R, Priefer U, Pühler A (1983) A broad host range mobilization system for *in vivo* genetic engineering: transposon mutagenesis in Gram negative bacteria. *Nat Biotechnol* 1: 784-791

Traglia GM, Chua K, Centron D, Tolmasky ME, Ramirez MS (2014) Whole-genome sequence analysis of the naturally competent *Acinetobacter baumannii* clinical isolate A118. *Genome Biol Evol* 6: 2235-2239

Tucker AT, Nowicki EM, Boll JM, Knauf GA, Burdis NC, Trent MS, Davies BW (2014) Defining gene-phenotype relationships in *Acinetobacter baumannii* through one-step chromosomal gene inactivation. *mBio* 5: e01313-01314

Wilharm G, Skiebe E, Higgins PG, Poppel MT, Blaschke U, Leser S, Heider C, Heindorf M, Brauner P, Jackel U *et al* (2017) Relatedness of wildlife and livestock avian isolates of the nosocomial pathogen *Acinetobacter baumannii* to lineages spread in hospitals worldwide. *Environ Microbiol* 19: 4349-4364
